# Human vulnerability to cancer malignancy is enhanced by evolution of higher mesenchymal CD44 expression compared to other mammals

**DOI:** 10.1101/2020.08.03.234617

**Authors:** Xinghong Ma, Anasuya Dighe, Jamie Maziarz, Edwin Neumann, Eric Erkenbrack, Yuan-Yuan Hei, Yansheng Liu, Yasir Suhail, Kshitiz, Irene Pak, Andre Levchenko, Günter P. Wagner

## Abstract

CD44 is a membrane-bound extracellular matrix (ECM) receptor interacting, among others, with hyaluronic acid (HA) and osteopontin (OPN). Cancer progression and metastasis are greatly influenced by the cancer micro-environment, consisting of ECM, immune cells and cancer-associated fibroblasts (CAF). Recruitment of fibroblasts (FB) into the role as CAFs is caused by paracrine signals from the tumor, including TGFb1, PDGF and OPN. The effect of OPN on the transformation of FB into CAF is mediated by CD44. CD44 expression in human skin and endometrial stromal fibroblasts (SF and ESF, respectively) also enhances invasibility of stroma by trophoblast as well as cancer cells. Here we study the evolution of *CD44* expression in therian mammals in both SF as well as ESF and demonstrate that the human lineage has experienced a concerted evolutionary enhancement of CD44 expression in SF and ESF, correlating with an increase in human vulnerability to cancer malignancy. In both human and cattle (*Bos taurus*), the dominant isoforms are CD44s and CD44v10 with 9 and 10 exons, respectively. CD44s is an isoform strongly associated with malignancy. In humans, an additional isoform is expressed: HsaCD44-205 with 8 exons not found in cattle. We show that the concerted increase of *CD44* expression in SF and ESF is largely due to cis-regulatory effects in the proximal promoter of *CD44*. We identify a primate specific acquisition of CEBPB binding sites in the CD44 promoter. Recruitment of CEBPB into CD44 regulation explains almost 50% of the lineage-specific increased *CD44* expression in primate skin fibroblasts but is not necessary for high CD44 expression in ESF. All these results suggest that selective modulation of *CD44* expression in skin fibroblasts could attenuate the cancer-promoting effect of CAF recruitment in the skin with minimal side effects on other cell types. Additional experimental data is needed to explore this possibility.

## Introduction

Cancer metastasis is the major cause of cancer-related mortality compared to the direct effects of the primary tumor. Malignancy rates differ greatly between mammalian species, broadly correlated with placenta type (D’Souza and Wagner 2014, Wagner, Kshitiz et al. 2020), where animals with less placental invasion tend to be also less vulnerable to cancer malignancy. Cancer progression and malignancy is in part the result of an interaction between tumor cells and the cancer associated stroma consisting of immune cells, cancer associated fibroblasts (CAF) and extracellular matrix (ECM) (Bhowmick and Moses 2005). We hypothesized that species differences in malignancy rate may be related to differences in the ability of stromal fibroblasts to resist invasion, a hypothesis we call ELI, for Evolved Levels of Invasibility (Kshitiz, Afzal et al. 2019). This reasoning applies to both, placental (trophoblast) invasion into the uterine endometrium as well as cancer malignancy, as variation in placental invasiveness in eutherian mammals is related to the ability of the uterine stroma to resist invasion (Samuel and Perry 1972). In a previous study we have shown that bovine skin and endometrial fibroblasts (SF and ESF respectively) are less invasible by trophoblast and melanoma cells than their human counterparts and that a knockdown of *CD44* in human cells decreases their invasibility, i.e. increases their resistance to trophoblast and cancer invasion (Kshitiz, Afzal et al. 2019). Here we document the evolution of *CD44* expression in mammals from the boreoeutherian clade of placental mammals and investigate the genetic basis of expression differences between human and bovines, i.e. the contributions of cis- and trans-effects to species differences in *CD44* expression. The importance of identifying the relative contribution of cis and trans-effects is that it has implications for the possibility of cell type specific modulation of *CD44* expression to attenuate the risk of malignant cancer spread.

CD44 is a well-known membrane bound receptor for ECM components that plays a major role in cancer progression and metastasis (Senbanjo and Chellaiah 2017, Chen, Zhao et al. 2018). CD44 expression in stromal fibroblasts is also important for the transformation of tissue fibroblasts into cancer-associated fibroblasts (CAF), which aid the tumor by supporting vascularization, ECM remodeling and travel with tumor cells to secondary cancer sites. The transformation of FB to CAF is caused by paracrine signaling from the primary tumor through, including but not limited to, TGFb1, PDGF and osteopontin (OPN, aka SPP1) (Sharon, Raz et al. 2015, Sahai, Astsaturov et al. 2020). Sharon and coworkers have shown that breast cancer cells secrete copious amounts of OPN and that tissue fibroblasts get transformed to CAF via the interaction of their CD44 and integrin avb3 receptors (Sharon, Raz et al. 2015). These two receptors play partially overlapping roles in CAF transformation, where integrin causes primarily the migratory phenotype of CAF and CD44 their pro-inflammatory effects. The critical role for malignancy of stromal CD44 is further supported by a study that shows that mesenchymal stem cells require CD44 to be transformed to CAF in a conditioned media assay as well as *in vivo* recruitment to a tumor site (Spaeth, Labaff et al. 2013). In addition, the different isoforms of CD44 have different potential to affect malignancy rate, with the CD44s isoform as the most associated with malignancy (Prochazka, Tesarik et al. 2014). Species differences in CD44 isoform expression in stromal cells can thus be a factor influencing the metastatic vulnerability of species. In our previous study we noted that human SF and ESF express higher levels of CD44 mRNA than cow and the knockdown of CD44 transcripts in human cells make human cells less invasible by cancer and trophoblast cells (Kshitiz, Afzal et al. 2019).

Here we show that high *CD44* expression in SF and ESF has specifically evolved in the primate lineage and is regulated by the homologous proximal cis-regulatory elements (CRE) in both cell types and species. Species differences in *CD44* expression are caused by both cis-as well as trans-regulatory factors. A statistical interaction effect between species and cell type suggests that part of the trans-acting factors are cell type-specific. We also identify CEBPB as a SF specific trans-factor explaining part of the higher *CD44* expression in human SF. These results suggest that cell type-specific modulation of CD44 expression and thus a tissue-specific suppression of CAF recruitment may be possible.

## Results and Discussion

### Humans evolved high expression of CD44 in stromal cells

To investigate gene expression evolution in skin fibroblasts (SF) and endometrial stromal fibroblasts (ESF) we cultured these two cell types from ten mammalian species, including the marsupial opossum (*Monodelphis domestica*) representing an outgroup to placental mammals, and nine species from the Boreoeutheria clade (roughly primate and rodent as well as ungulates and carnivores) of placental (eutherian) mammals (Figure 1A). Gene expression was assessed by bulk RNAseq of isolated cultured fibroblast and quantified as transcripts per million transcripts (Li, Sun et al. 2009, Wagner, Kin et al. 2012) based on the 8,639 one-to-one orthologous genes among these 10 species. Each gene was quantified using species-specific transcript reconstructions from the read data (see M&M). Here we focus on the results for CD44. A more comprehensive analysis of this data will be presented elsewhere.

**Figure 1:**
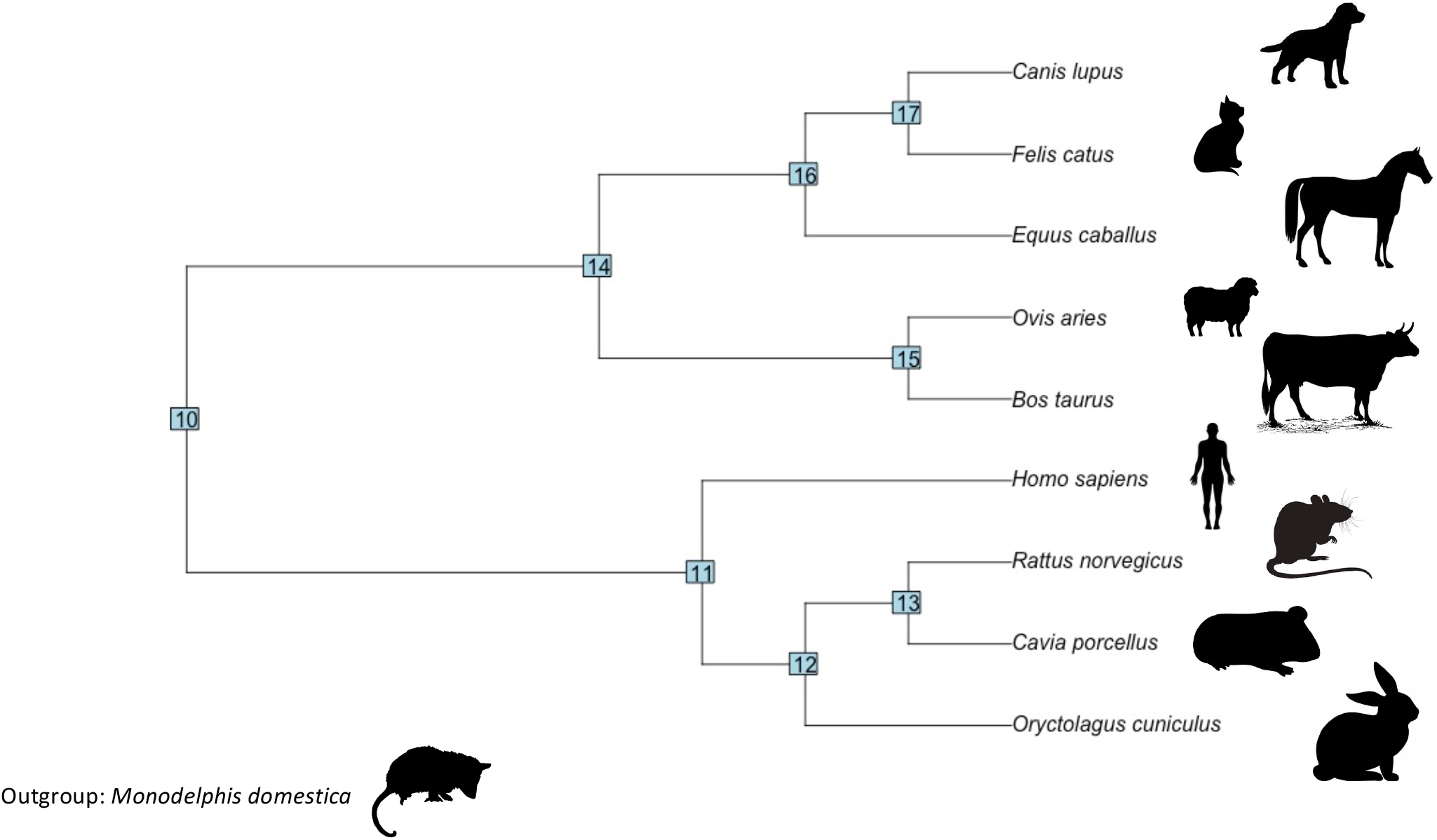
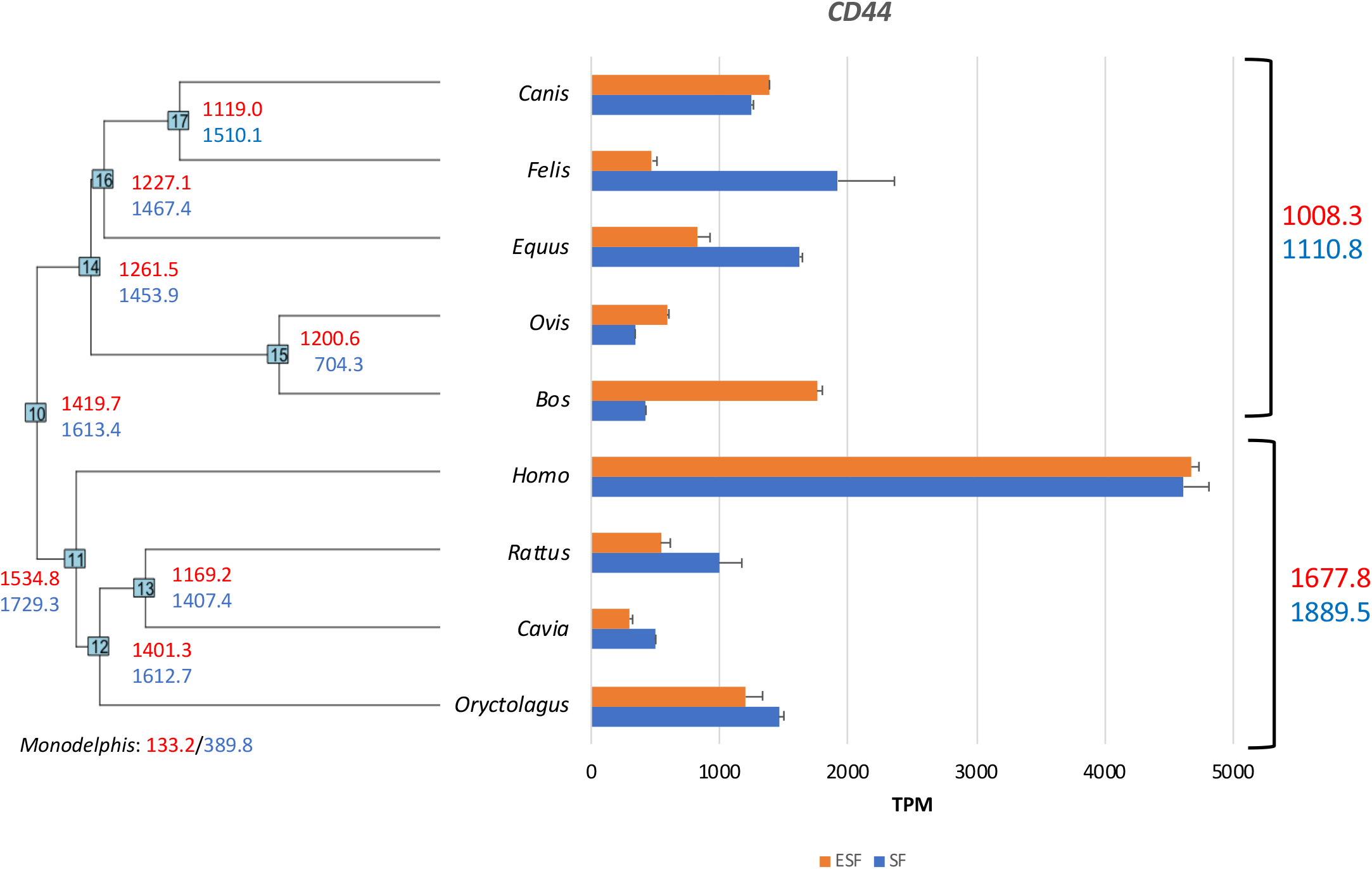
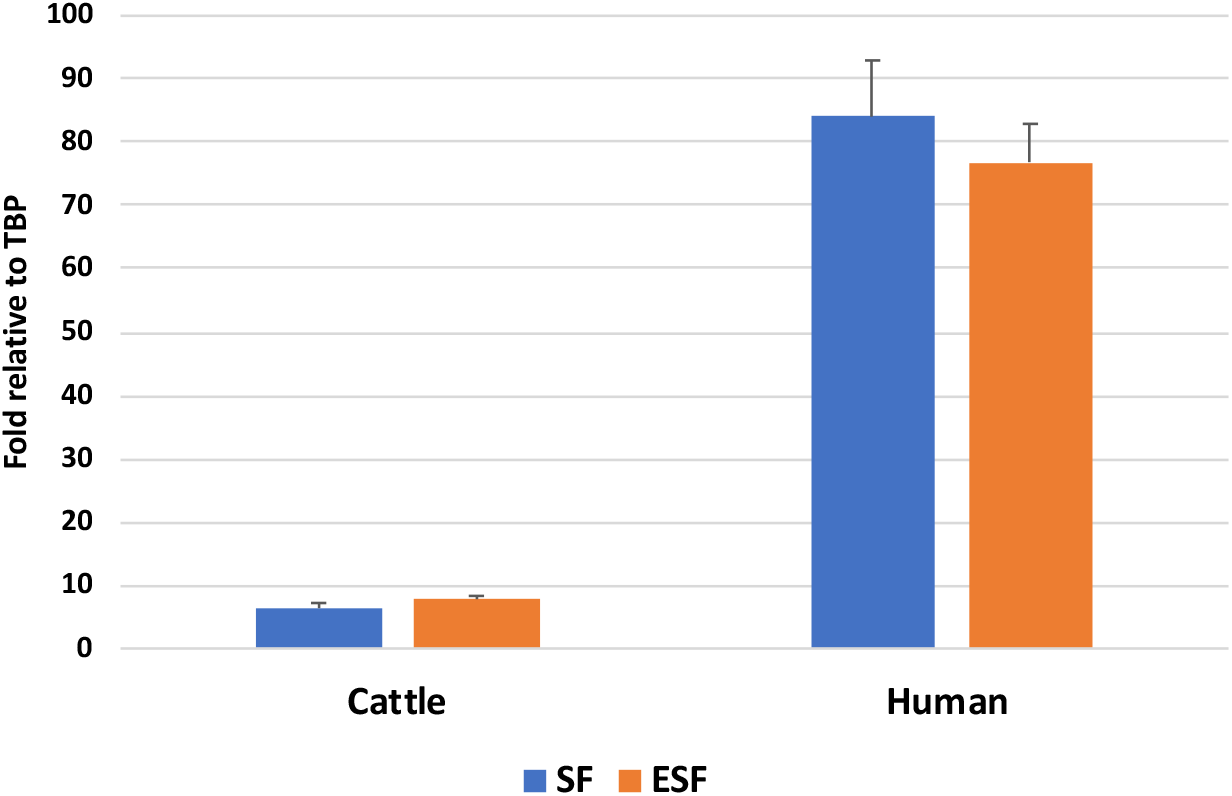
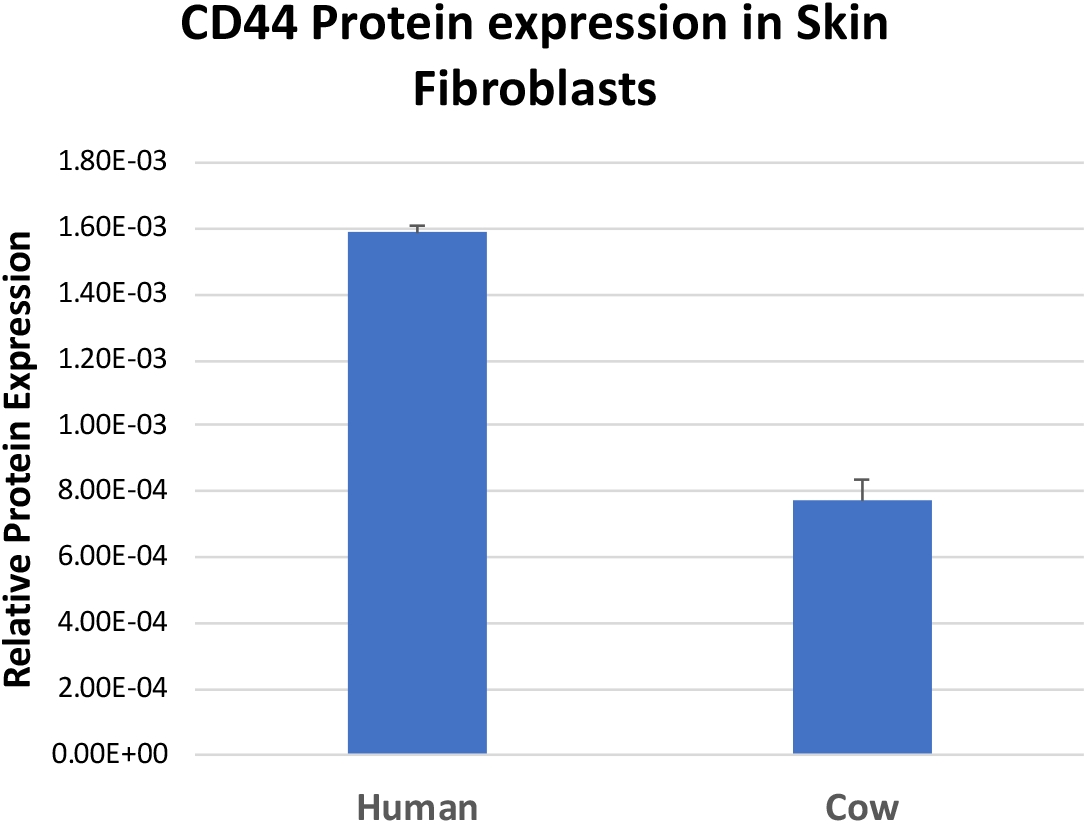
CD44 expression in Boreoeutherian mammals. A) Taxon sample with phylogenetic relationships among the species. B) Gene expression levels [TPM] in skin and endometrial fibroblasts from nine Boreotherian mammals and the opossum *Monodelphis domestica* as an outgroup. At the nodes of the tree are the ancestral state reconstructions based on REML algorithm (see M&M). Red numbers are TPM in endometrial stromal firbroblasts (ESF) and blue numbers are for skin fibroblasts (SF). C) qPCR confirmation of the high expression of *CD44* in human in comparison to cow. D) CD44 protein expression in skin fibroblasts from human and cow. Human skin fibroblasts express at least twice as much CD44 protein than cow skin fibroblasts.

Figure 1B shows the expression levels of CD44 mRNA in both SF and ESF for the ten species included in this study. At the nodes of this tree ancestral gene expression estimates are noted using Residual Maximum Likelihood (REML) method (Schluter, Price et al. 1997, Felsenstein 2003). As can be seen from these data, the expression in all animals sampled, excluding humans, is below 2,000 TPM (average w/o human 1062 TPM for SF and 885 TPM for ESF) and does not show very distinct evolutionary trends. However, in both human ESF and SF the expression level is between 4,600 and 4,700 TPM. We confirmed this large discrepancy in CD44 expression between human and bovine cells with qPCR (Figure 1C).

Interestingly, opossum has about a 10-fold lower expression level of *CD44* than even the nonhuman placental mammals, 133 TPM and 390 TPM in ESF and SF, respectively, suggesting an increase in expression associated with the evolution of invasive placentation which happened in the stem lineage of eutherian mammals (Mess and Carter 2006, Wildman, Chen et al. 2006, Elliot and Crespi 2009).

Phylogenetic reconstruction of ancestral gene expression levels suggests that the common ancestor of Euarchontoglires had an expression level of about 1,535 TPM in ESF and 1,729 TPM in SF, which suggests a threefold increase of *CD44* expression in the primate lineage. For SF, for which we have also data from *Macaca mulatta* we find that the level of gene expression in monkey SF is statistically indistinguishable from that in human SF. This suggests that the high expression of *CD44* in human skin fibroblasts evolved before the most recent common ancestor of Catarrhini (old world monkeys and apes) and after the ancestor of Euarchontoglires (primates, rodents and rabbits).

### CD44 protein abundance in human and bovine skin fibroblasts

In order to investigate whether the RNA abundance differences between human and bovine are indicative of corresponding protein abundance differences we investigated the relative expression level of CD44 proteins in human and cow skin fibroblasts. Since protein quantification across species using antibodies is problematic due to species differences in the amino acid sequences of the target protein which can cause differences in reactivity with the antibody, we resorted to a new and reproducible quantitative mass spectrometry (MS) technique (See Methods). We compared the ratio of CD44 MS quantities from each sample to the total protein signals in the MS run. The reference set of proteins for each sample was the overlap between the proteins detected in human and bovine samples (6,652 and 5,510 proteins for human and cow, respectively, with an overlap of 4,269) to assure commensurable expression scales for human and cow samples (Figure 1D). The relative CD44 protein abundance in bovine skin fibroblasts was lower than in human samples, as expected (t-test p=7.25 10^−3^). The fold difference of human compared to cow was 2.05x, which is smaller than the fold differences based on RNAseq and qPCR (10.9x and 13.1x respectively). Whether this differences in RNA vs protein abundances is due to regulation at the level of translation, protein stability or due to post-translational modifications affecting peptide quantification by MS is not clear.

### CD44 splice isoform expression

*CD44* is known to express a large number of splice isoforms which play different biological roles (Prochazka, Tesarik et al. 2014, Senbanjo and Chellaiah 2017, Chen, Zhao et al. 2018). Here we investigated the expression of isoforms from our read data in ESF and SF of human and cow.

In humans the CD44 gene has been described as having 19 exons divided in 9 constant exons 1, …,8 and 10 (“exon” 9 never expressed) and nine so-called variable exons, called v2 to v10 (Screaton, Bell et al. 1992). In quantifying transcript abundance we focused on high quality protein coding transcript annotations that correspond to consensus coding sequences (CCDS) and we refer to them using the ENSEMBL transcript names with the format HsaCD44-2xx. More than 96% of transcripts belong to three isoforms: HsaCD44-201, HsaCD44-210 and HsaCD44-205 (Figure 2A), where the far most dominant isoform is HsaCD44-201 with 9 exons and 361 amino acids (Figure 2B & C), which in the literature is sometimes called the standard isoform CD44s. HsaCD44-210 is identical to HsaCD44-201 with the addition of the most 3’ of the so-called variable exons, ENSE00003608645=v10, and can thus be called CD44v10 following a naming tradition in the biomedical literature (Figure 2A). Not much is known about the functional roles of this specific isoform. The third transcript, HsaCD44-205, is unusual as it has 8 exons and is lacking one of the so-called “constant” exons of HsaCD44-201/CD44s, namely ENSE00003526469 (Figure 2A and Suppl. Figure 1A). Differences in annotated 3’ and 5’ exons are not affecting the predicted amino acid sequence (Suppl. Figure 1B).

**Figure 2:**
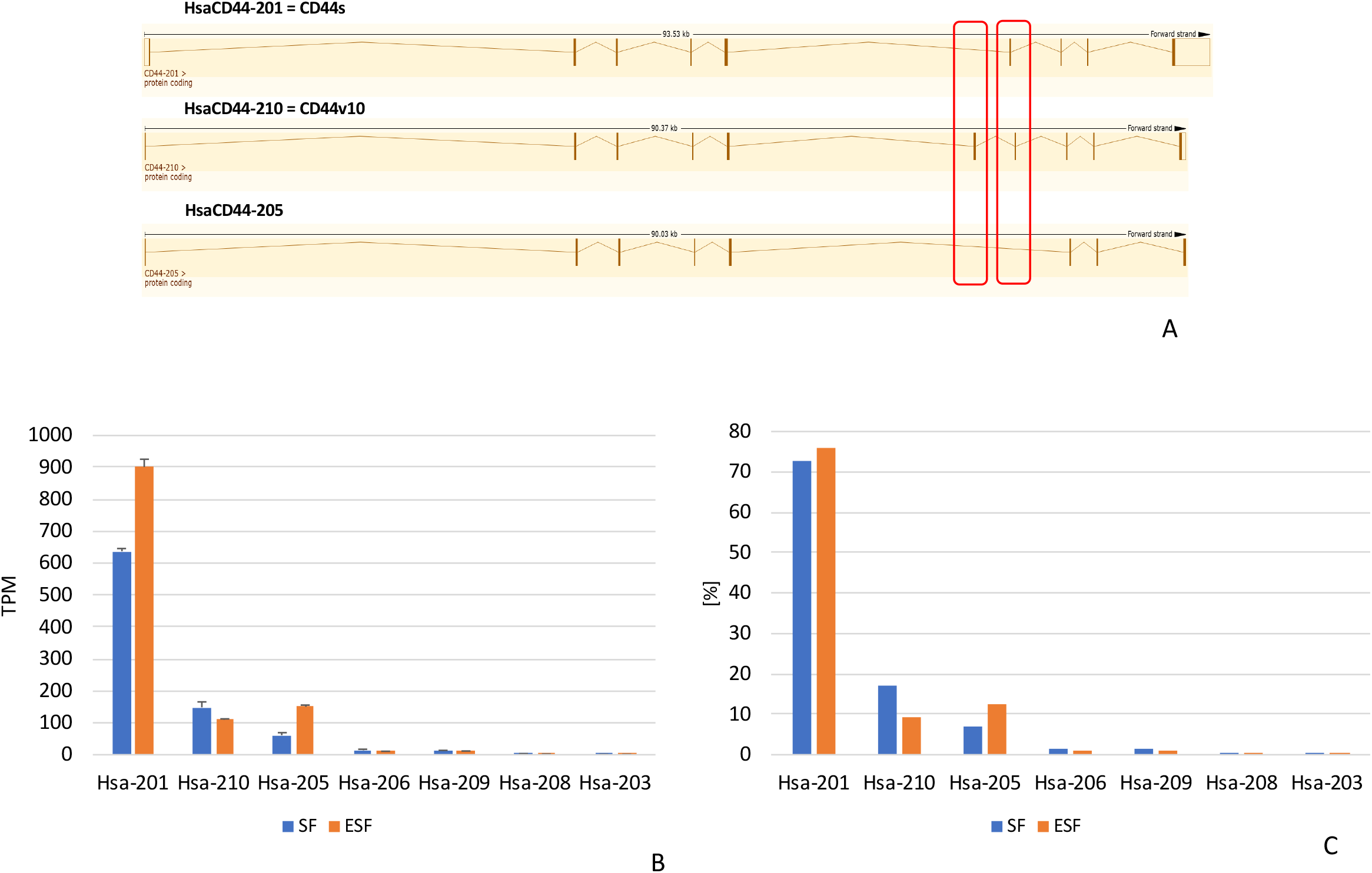
CD44 RNA isoform expression in human skin and endometrial fibroblasts. A) intron-exon structure of the three dominant isoforms from human mesenchymal cells: HsaCD44-201 which is also known as CD44s, HsaCD44-210 which retains the variable exon 10, Hsav10, and HsaCD44-205 which lacks one of the “constant” exons. [images courtesy of ENSEMBLE.org] B) expression levels [TPM] of isoforms in SF and ESF. Note that only three isoforms have considerable levels of expression. C) relative expression levels [%] of CD44 isoforms in human skin and endometrial fibroblasts.

In bovine cells the majority of transcripts belong to two isoforms, BtaCD44-209 and BtaCD44-208, which together make up >93% of transcripts in ESF and 98.9% in SF (Figure 3A & B and Suppl. Figure 1C). In both cell types the dominant transcript is BtaCD44-209 with 79% and 87% representation, respectively (Figure 3C). Note that the ENSEMBLE transcript numbers in different species are not indicating homology. Inspection of the annotated sequences revealed that BtaCD44-209 is homologous to HsaCD44-201 and is thus a bovine homolog of human CD44s (Suppl.Figure 2), and BtaCD44-208 is homologous to HsaCD44-210 (Suppl. Figure 3), which we call CD44v10.

In both species and cell types the most abundant protein coding transcript is CD44s (HsaCD44-201 and BaCD44-209), with a minor contribution of CD44v10 (HsaCD44-210 and BtaCD44-208). In addition, human cells express a smaller isoform of 8 exons, HsaCD44-205, which is not much discussed in the literature. Hence human stromal cells differ from cow stromal cells by both an overall higher expression of CD44 (Figure 1B) and the presence of a minor isoform, HsaCD44-205 (Figures 2 and 3) while the relative abundance of the two major isoforms is the same in skin and endometrial stromal fibroblasts in both species.

**Figure 3:**
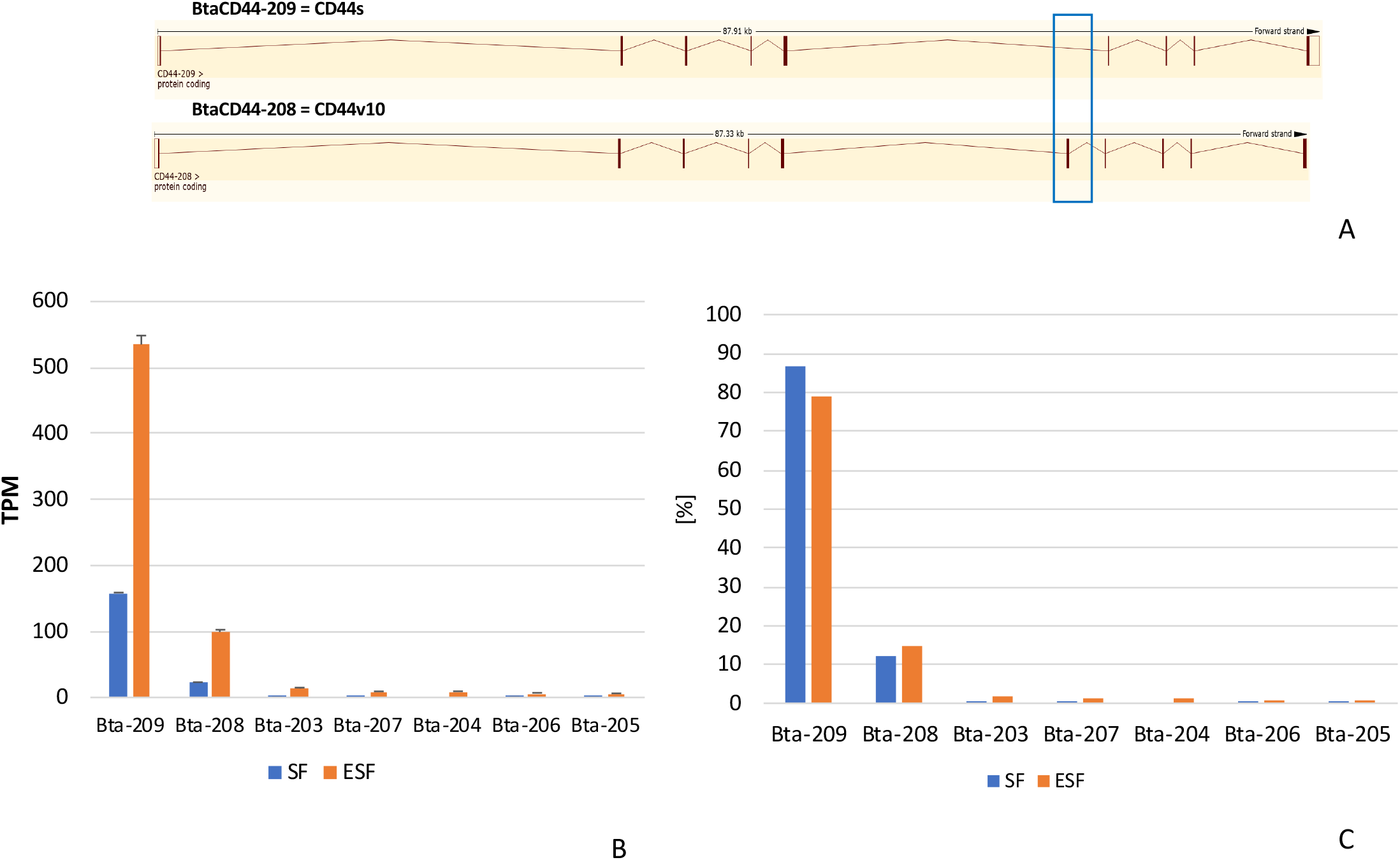
*CD44* RNA isoform expression in bovine skin and endometrial fibroblasts. A) intron-exon structure of the two dominant isoforms from bovine mesenchymal cells: BtaCD44-209 which is also known as CD44s, BtaCD44-208 which retains the variable exon 10 like the human HsaCD44-210 (see Figure 2A). [images courtesy of ENSEMBLE.org] B) expression levels [TPM] of isoforms in SF and ESF. Note that only three isoforms have considerable levels of expression. C) relative expression levels [%] of CD44 isoforms in bovine skin and endometrial fibroblasts.

### CD44 *is transcribed from the same promoter in skin- and endometrial fibroblasts*

The RNAseq and qPCR results suggest that the primate lineage saw an increase of *CD44* RNA expression in both cell types, SF and ESF (Figs. 1B and 1C). A concerted change in gene expression can be explained either by the fixation of pleiotropic mutations that affect gene expression in both cell types (Musser and Wagner 2015, Liang, Musser et al. 2018) or by natural selection acting on gene expression in both celltypes even though the genetic variation may be uncorrelated. On the mechanistic level, pleiotropic change in gene expression can either be due to mutations in cis-regulatory elements (CRE) for CD44 that are active in both cell types, or changes in the activity of trans-factors that affect CD44 expression in both cell types. We investigate these two possibilities (cis- and trans-regulation) in this and the next sections.

To investigate whether the evolution of *CD44* gene expression can be due to changes in the same cis-regulatory element or by species specific promoters, we first asked whether the *CD44* RNA was transcribed from the same promoter in both cell types and species. We performed a 5’RACE experiment on cDNA from both cell types and species and found that the 5’RACE fragments recovered are of similar length (~460bp, with about a ~120bp 5’UTR) (Suppl Figures 4A and 5B) and amplify orthologous sequences (Suppl. Figure 4B). We note that the 5’UTRs reported in annotated transcript sequences in ENSEMBLE differ greatly in length from the 5’ UTR that we amplified (see above). For instance, the 5’UTR of HsaCD44-201 is reported to be 434bp. We could not trace the tissue source for this cDNA from where the HsaCD44-201 was isolated, but the different results suggest that the cDNA was cloned from another cell type than those investigated here, suggesting that *CD44* in cell types other than skin and endometrial fibroblasts are controlled by an alternative promoter.

From these results we concluded that the promoters used in the two mesenchymal cell types, SF and ESF, are the same and different from other cell types. Thus it is possible that the proximal CRE shared between SF and ESF can be responsible for the concerted increase in CD44 expression in the human fibroblasts. We tested this possibility with reporter gene experiments.

### Reporter gene expression recapitulates species and cell type differences

In order to measure the contribution of cis and trans factors to the gene expression difference between cow and human ~3kb fragments were cloned upstream from the TSS from the bovine and human *CD44* locus’ TSS into a reporter construct (see M&M). We call these fragments pCRE for “proximal Cis-Regulatory Element”. We compared the reporter gene RNA expression driven by the human and cow pCRE in their cognate cell types (Figure 4A). The activity of the human pCRE in human ESF and SF is higher than the cow pCRE in cow cells (fold differences for ESF and SF: 3.0x and 6.7x respectively; t-test of lnFOLD difference p=4.5 10^−4^ and p=7.5 10^−5^). Furthermore, the activity of the human pCRE in the two human cell types is statistically indistinguishable (p=0.671), but in the bovine cells the activity of the bovine pCRE is lower in SF than in ESF (2.4x; p= 1.8 10^−2^). We conclude that the activity of the reporter system qualitatively recapitulates the pattern seen in the intrinsic *CD44* RNA expression measured by RNAseq or qPCR among species and cell types (Figures 1B and 1C).

**Figure 4:**
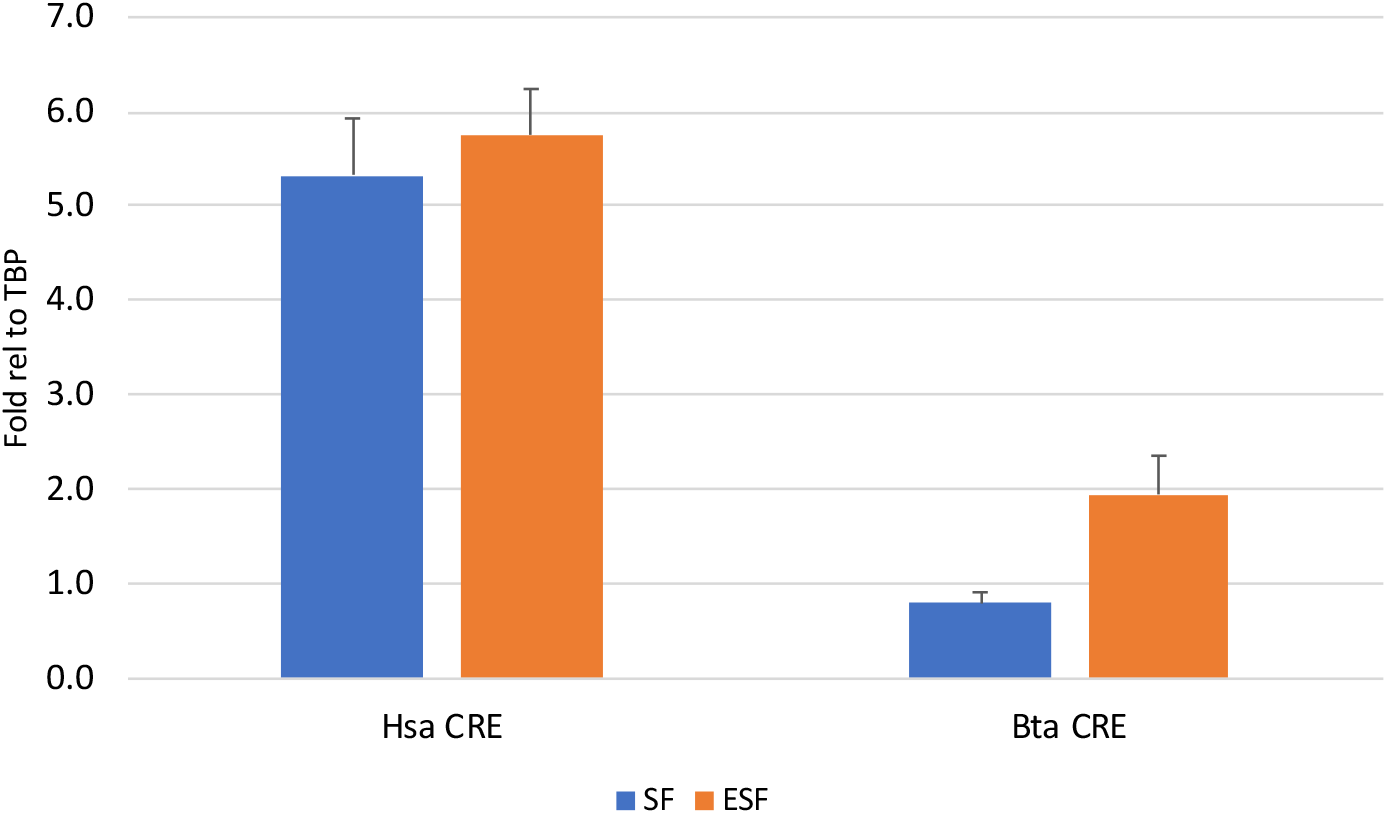
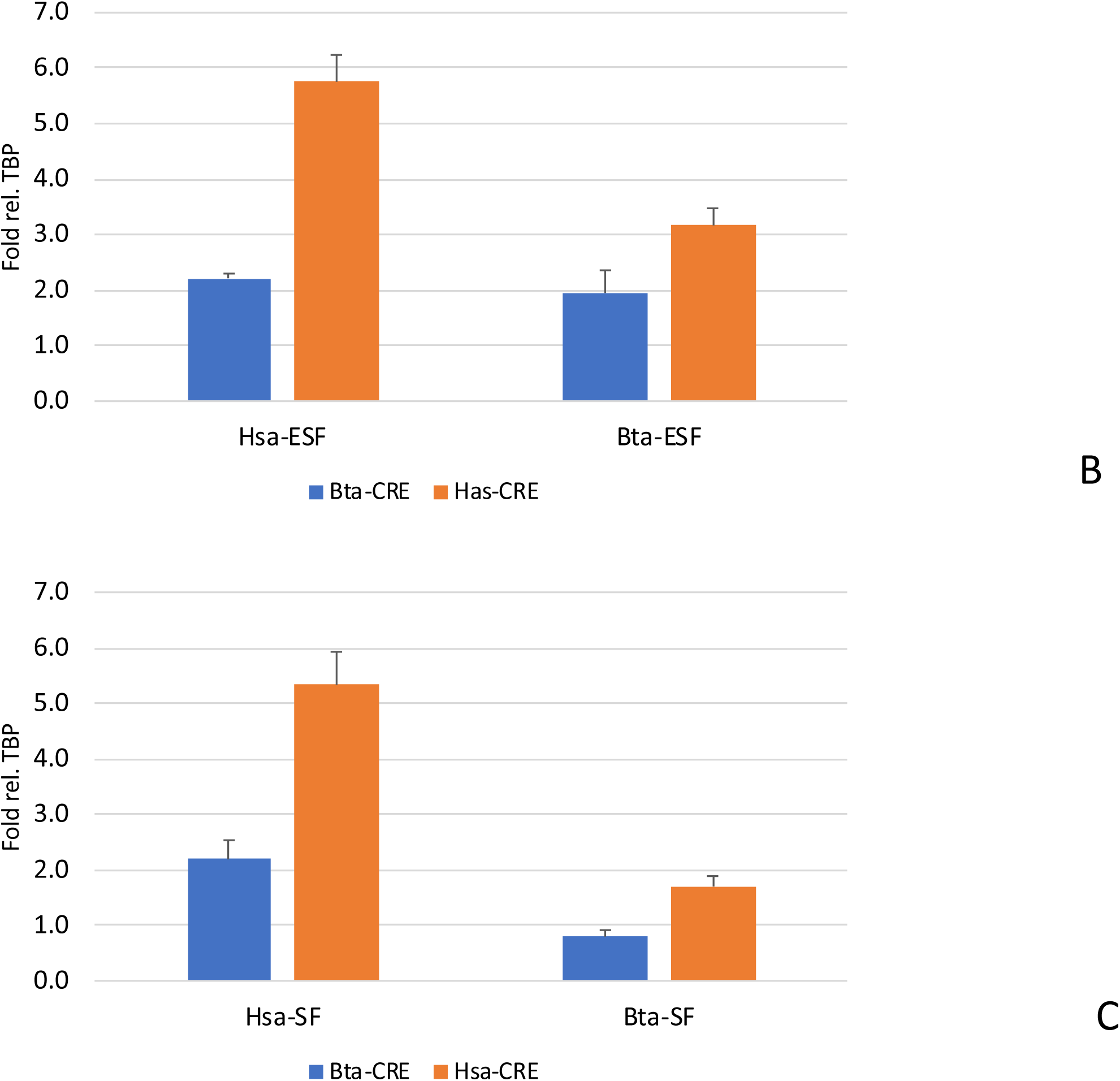
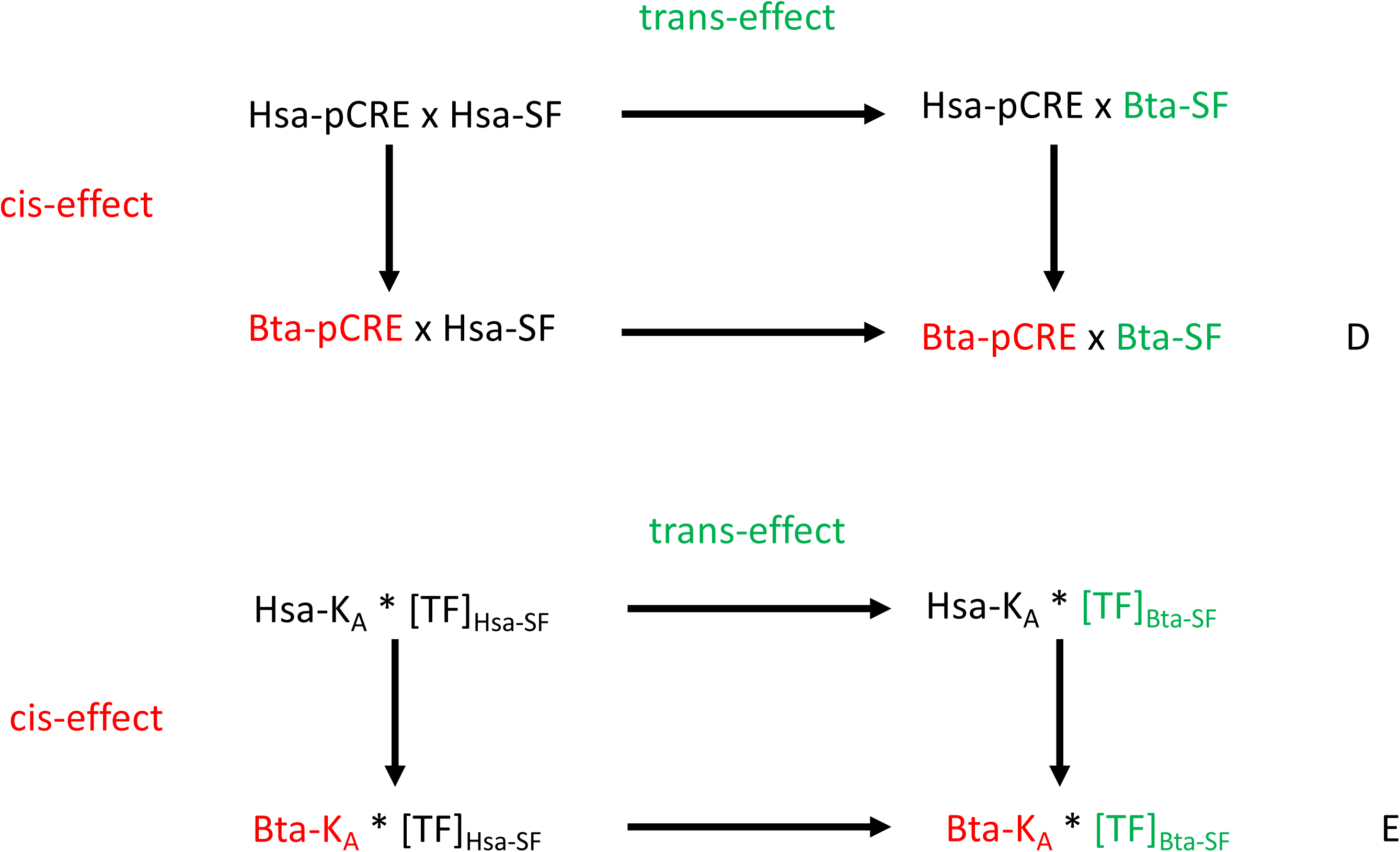
results of reporter gene expression experiments with human and bovine CD44 promoters in human and bovine skin and endometrial fibroblasts. A) expression of reporter gene transcripts [qPCR fold difference to TBP internal reference] of human and bovine promoters in their cognate cells, i.e. human promoter in human cells and bovine promoter in bovine cells. Note that the pattern of expression qualitatively reproduces the cell type and species typical expression levels found in RNAseq (see Figure 1B). B) Cross species comparison of promoter activity in skin fibroblast cells. Blue bars are results for bovine promoter (BtaCRE), red for human promoter (HsaCRE); left columns tested in human skin fibroblasts (Hs SF) and right columns tested in bovine skin fibroblasts (Bta SF). Note that in cells from both species the bovine promoter drives lower reporter gene expression than the human promoter supporting the interpretation that some of the differences in human and bovine gene expression are caused by differences in the promoter sequences. In addition the finding that both promoters drive a higher reporter gene expression in human cells also gives evidence for trans-regulatory differences between species contributing to the expression difference between cows and humans. C) Cross species comparison of promoter activity in endometrial fibroblasts. The annotation is analogous to B) and the results also suggest that both cis-as well as trans-regulatory factors contribute to the species differences in *CD44* expression. D) Experimental setup for quantifying cis- and trans-effects at the example of skin fibroblasts. Hsa-pCRE is the human promoter, Hsa-SF are human skin fibroblasts, Bta-pCRE is the bovine promoter, and Bta-SF are bovine skin fibroblasts. For instance, Hsa-cCRE x Hsa-SF means that the human promoter is tested in human skin fibroblasts etc.. Comparing fold differences along the arrows yields estimates of the corresponding cis and trans-regulatory effects. E) Predictions of the “ideal gene” theory (see Appendix) for the outcome of the experimental setup illustrated in D. Note that according to the model cis and trans-effect should combine multiplicatively and thus should not lead to statistical interaction effects as measured on the multiplicative scale. This is what the experimental results in fact show (see text for details).

### Both cis and trans-regulatory factors contribute to species and cell type differences

Next we consider the activities of the human and bovine pCRE in all combinations of cell types and species of origin (Figures 4B and C). A two-way ANOVA on the log transformed fold values for SF reveals strong evidence for both cis-as well as trans-effects (p=8.0 10^−5^, F=27.73, 1/16 dgf for the cis-effect, and p=6.25 10^−6^, F=43.39, 1/16 dgf for the species trans-effect among SF) with similar effect sizes (on average 2.28 fold cis-effect and 2.94 fold trans-effect). Similar results were found for ESF. The significance levels of cis- and trans-effects in ESF were 2.73 10^−6^ (F=34.24, 1/28 dgf) and 1.88 10^−3^ (F=11.77, 1/28 dgf) respectively. An interesting detail is that the size of the cis-effect is comparable in both cell types (2.28x for SF and 2.05x for ESF), while the trans-effects are different by a factor 2 (2.94x in SF and 1.44x in ESF) hinting to cell type specific trans-regulatory landscapes.

An unexpected result was that in both cell types there was no interaction effect between cis and trans-regulatory factors in spite of strong direct effects. This is surprising given the complex molecular mechanisms underlying eukaryotic transcriptional regulation which intuitively should lead to non-linearities and thus to statistical interaction effects. To appreciate this result we briefly remind our readers of the scale dependence of interaction effects (Wagner 2015)(see also Appendix). The response variable are qPCR data that exists on a multiplicative scale, i.e. fold change, with respect to an internal standard (TATA binding protein, *TBP*). For that reason tests for interaction have to test for interactions at the multiplicative scale (Wagner 2010, Wagner 2015). However, the ANOVA model treats effects on an additive scale, and therefore the ANOVA test has to be performed on a log-fold change scale. In fact, the result reported above was obtained by ANOVA on the log-fold change. The same test performed on the original multiplicative scale suggests a mild interaction effect, but this is an artifact due to inappropriate scale choice.

To interpret the result we analyzed a simple kinetic model of transcriptional regulation (see Appendix), where the rate of production of mRNA is proportional with the occupancy frequency of a transcription factor at a CRE. The analysis predicts that the equilibrium reporter gene concentration 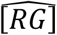 is,

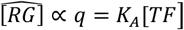

if q<<1. Applying this model to the experimental outline (Figure 4D) predicts that, on the multiplicative scale there should be no interaction effects between the trans-factor (Figure 4E), i.e. the transcription factor concentration [*TF*], and the cis-factor, i.e. the binding constant to the CRE, *K_A_*. Of course, the consistency between our experimental results and the model does not mean that the *CD44* promoter is regulated by one transcription factor, even in our artificial reporter gene assay. What it does mean, however, is that whatever differences exist in the trans-regulatory environment in human and cow cells and *CD44* promoters their differences are dynamically equivalent to the effect of a single trans-factor. The same result is obtained in a two transcription factor model if their influence on gene transcription is due to a cooperative effect (Appendix), also known as an “AND” gate, meaning that the rate of transcription is dominated by the co-occupancy of the CRE by both transcription factors. In contrast, a model where the two transcription factors influence transcription independently is predicted to cause interaction effects (Appendix). We conclude that our finding of no detectable interaction among cis- and trans-factors (species origin of promoters and cells) on the multiplicative scale is consistent with a model where either a single transcription factor or a set of cooperative transcription factors are responsible for the trans-effects.

We next performed a three way ANOVA with these three factors: species of cell origin, cell type, SF or ESF, and species of promoter origin with log-Fold change as response variable (Table 1). The results provide strong evidence for species, cell type and cis-regulatory effects, like the separate two-way ANOVAs for each cell type. The p-values for all the direct effects is between 10^−10^ and 10^−3^. In addition there is a significant interaction effect between species and cell type (p= 4.7 10^−3^; F=8.87, 1/44 dgf). To interpret this result we again turned to the kinetic model of transcriptional regulation.

**Table 1:** three way ANOVA of the reporter gene experiments with human and cow pCRE in human and cow skin and endometrial fibroblasts. The response variable is the log transformed fold difference measured by qPCR.

| *Source of Variation* | *SS* | *d.f.* | *MS* | *F* | *p-level* |
| --- | --- | --- | --- | --- | --- |
| *Factor #1 (species of cell)* | 6.21209 | 1 | 6.212 | 45.60 | 2.64E-08 |
| *Factor #2 (promoter)* | 8.39617 | 1 | 8.396 | 61.63 | 6.62E-10 |
| *Factor #3 (cell type)* | 1.88854 | 1 | 1.889 | 13.86 | 5.60E-04 |
| *Factor #1 + #2 (species of cell x promoter)* | 0.2093 | 1 | 0.209 | 1.54 | 2.22E-01 |
| *Factor #1 + #3 (species of cell x cell type)* | 1.2089 | 1 | 1.209 | 8.87 | 4.69E-03 |
| *Factor #2 + #3 (promoter x cell type)* | 0.02672 | 1 | 0.027 | 0.20 | 6.60E-01 |
| *Factor #1 + #2 + #3 (species of cell x promoter x cell type)* | 0.02914 | 1 | 0.029 | 0.21 | 6.46E-01 |
| *Within Groups* | 5.99452 | 44 | 0.136 |  |  |
| *Total* | 23.96538 | 51 | 0.470 |  |  |

Analyzing the most parsimonious model of gene regulation for an experiment assessing species and cell type effects shows that absence of an interaction effect would require quite special conditions (Appendix). Specifically the model shows that interaction effects to be absent would require that the ratio of expression levels of the upstream regulators between cell types has to be that same for both species:

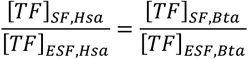

Conversely, if one finds interaction effects between species origin of cell and cell type, then there are species specific trans-regulatory differences between cell types. The fact that we find such interaction effects suggest that the trans-regulatory factors for *CD44* expression evolves quasi-independently among cell types, which confirms the idea that SF and ESF are in fact different cell types (Arendt, Musser et al. 2016).

### *Regulation of* CD44 *in mesenchymal cells*

A recent review of the role of *CD44* in cancer biology has summarized the current knowledge about upstream regulatory factors for *CD44* expression in cancer cells (Chen, Zhao et al. 2018). Positive regulators are SP1, EGR1, TCF4, AP-1, NSFKB and ETS-1. Negative regulators are p53, KLF4 and FOXP3. In order to find an explanation for the high *CD44* expression in human mesenchymal cells we looked for positive regulators that are more expressed in human cells and negative regulators that are more expressed in bovine cells (Suppl. Figure 6). The positive transcription factors with human biased expression are *SP1, TCF4* and *NFKB* (Suppl. Figure 6A). We tested all three of them with siRNA mediated knockdown experiments in human ESF and did not detect any change in *CD44* expression (Suppl.Figure 7A). Only one transcription factor among the negative regulators has a bovine-biased expression, *KLF4* (Suppl. Figure 6B). We tested the effect of KD of KLF4 in cow SF and ESF and could not detect any upregulation of CD44 in these cells (Suppl. Figure 7B). We conclude that the regulators identified in the cancer literature, summarized in Chen et al. (2018), do not include the trans-regulatory factors contributing to the high expression of *CD44* in human mesenchymal cells.

To find cis regulatory candidates that could explain the high *CD44* expression in human mesenchymal cells we mapped transcription factor binding sites in the upstream 5 kb of the CD44 locus from human, rabbit, guinea pig, rat, horse, sheep and cow and identified transcription factors with binding site abundance in human higher than any of the other species and expression levels in both human ESF and SF of >>3TPM (Table 2). This search revealed six candidate transcription factors (Suppl. Figure 8). These are candidate transcription factors that could explain the cis-regulatory effects on CD44 expression. Among them the top candidate is CEBPB, a well-known transcription factor essential for the decidualization of human ESF and other mesenchymal cell types such as adipocytes. We identified three transcription factor binding site in the human *CD44* locus, and the RNA expression of CEBPB is about 95 TPM in both human cell types.

**Table 2:**
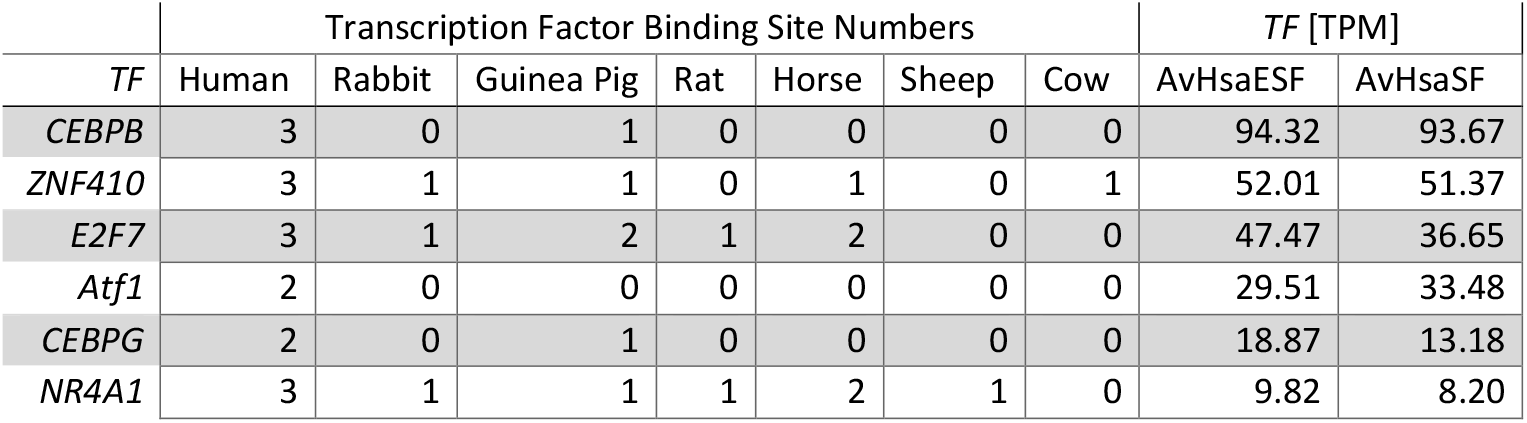
Transcription factor binding site numbers at the CD44 promoter region as well as RNA expression levels of the corresponding transcription factor in human mesenchymal cells in TPM. Note that CEBPB has the most consistent difference in terms of binding site numbers of human relative to other species and the highest expression level in human cells and is thus the strongest candidate to explain part of the lineage specific expression levels of *CD44*.

| *TF* | Transcription Factor Binding Site Numbers |  |  |  |  |  |  | *TF* [TPM] |  |
| --- | --- | --- | --- | --- | --- | --- | --- | --- | --- |
|  | Human | Rabbit | Guinea Pig | Rat | Horse | Sheep | Cow | AvHsaESF | AvHsaSF |
| *CEBPB* | 3 | 0 | 1 | 0 | 0 | 0 | 0 | 94.32 | 93.67 |
| *ZNF410* | 3 | 1 | 1 | 0 | 1 | 0 | 1 | 52.01 | 51.37 |
| *E2F7* | 3 | 1 | 2 | 1 | 2 | 0 | 0 | 47.47 | 36.65 |
| *Atf1* | 2 | 0 | 0 | 0 | 0 | 0 | 0 | 29.51 | 33.48 |
| *CEBPG* | 2 | 0 | 1 | 0 | 0 | 0 | 0 | 18.87 | 13.18 |
| *NR4A1* | 3 | 1 | 1 | 1 | 2 | 1 | 0 | 9.82 | 8.20 |

In order to test whether the so identified putative primate specific trans-regulatory factors are causally relevant, we focused on CEBPB using a siRNA knockdown approach. The KD efficiency was high, at about 90% reduction on average; experiments with lower KD efficiency were ignored. In human ESF the CEBPB KD did not affect *CD44* expression, but in SF we found a consistent reduction of about 30% of *CD44* expression at days one and two after KD but no effect on day three (Figure 5A). We interpret these results as indicating that CEBPB is relevant for *CD44* expression in human skin fibroblasts but not in human ESF, consistent with the finding above that ESF and SF diverged in their trans-regulatory landscape (see section above). In addition, the results suggest that CEBPB has a cumulative (so-called “additive”, although not technically *additive*) effect on *CD44* expression in human skin fibroblasts. Note that the role of CEBPB in regulating *CD44* transcription is likely specific to the primate lineage, given the lack of CEBPB binding sites in the *CD44* promoter region of other species. Hence CEBPB and other putative primate specific trans-factors are added on to the ancestral gene regulatory network for CD44 regulation late in evolution and are thus likely modulatory rather than essential. Further, the results suggest that the loss of CEBPB after KD is compensated by other factors over a time period of three days. All these results suggest that *CD44* trans-regulation is redundant and robust.

The influence of CEBPB on *CD44* expression accounts for about 30% of the expression in human skin fibroblasts. Given the distribution of transcription factor binding sites we suggest that CEBPB is in part responsible for the lineage specific increased CD44 expression in the primates. The lineage specific increase of CD44 expression is about 3x, and thus the recruitment of CEBPB explains about half of the increase in the primate lineage (Figure 5B).

**Figure 5:**
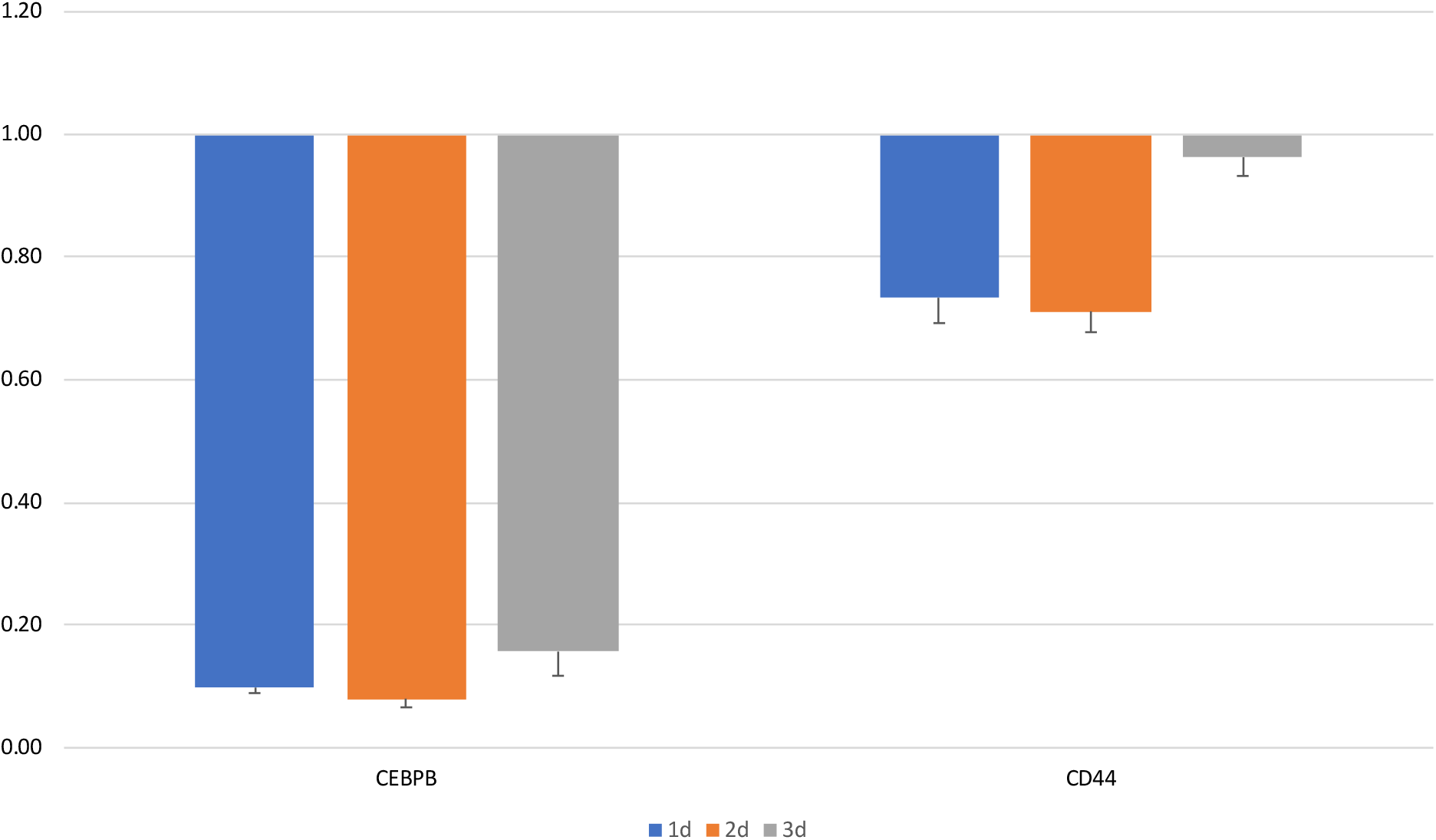
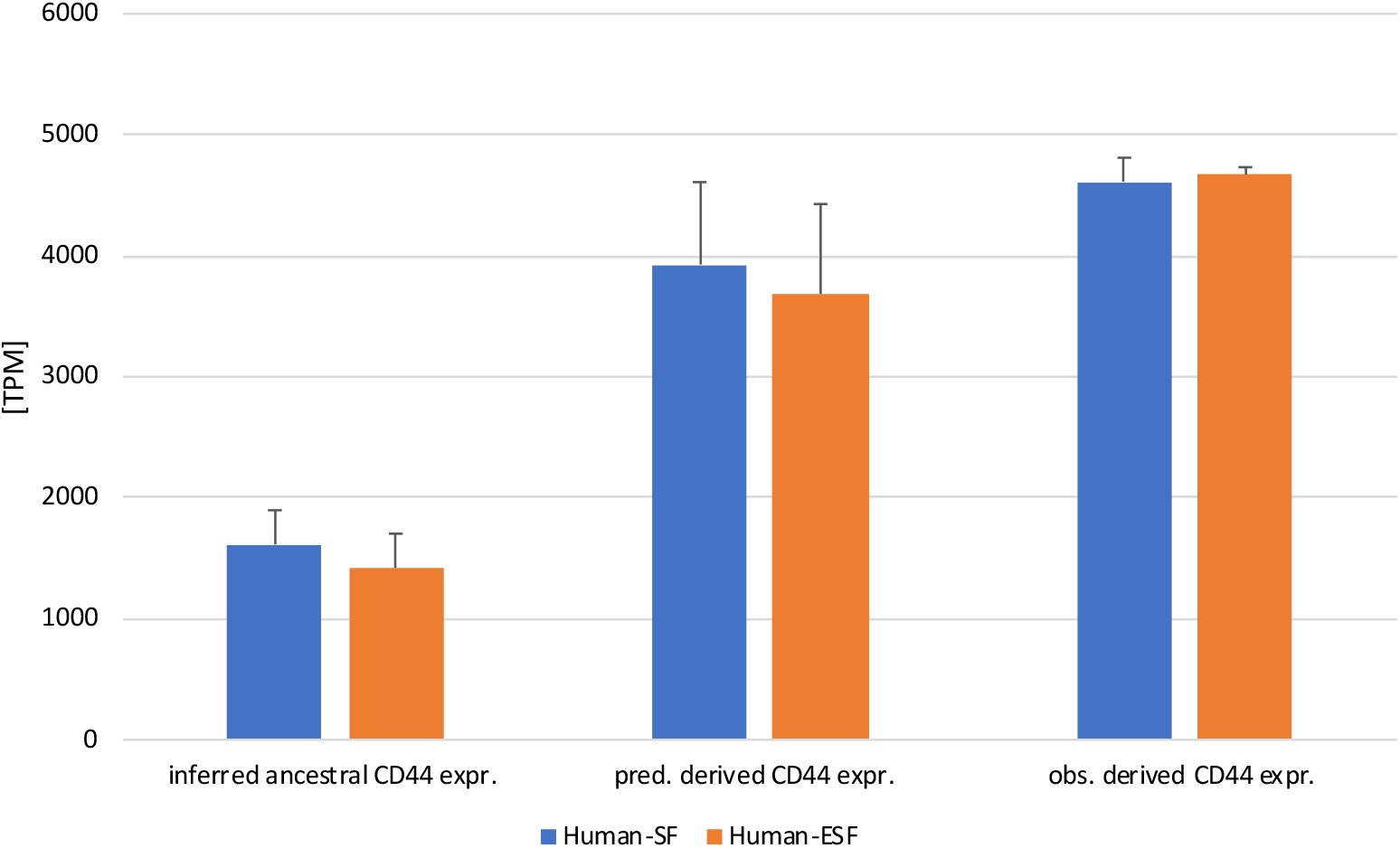
cis regulatory factors of CD44 expression in human fibroblasts. A) The impact of CEBPB knockdown (KD) on *CD44* expression in human skin fibroblasts after one (blue), two (red) and three days (gray) of knockdown. The KD efficiency is larger than 80% over all time periods. After one or two days of KD the expression level of CD44 is reduced by about 30% and after three days there is no effect detectable (likely because of compensation by other regulators). B) explanation, in the human lineage, of the evolutionary change in CD44 expression by cis-effects measured in the report gene assay. Left column, inferred CD44 expression [TPM] in the most recent common ancestor of Boreoeutheria (see Figure 1B). Middle column, predicted evolutionary outcome of CD44 expression explained by the cis-regulatory effects measured in reporter gene assay (Table 3, Suppl. Figure 10). Right column, measured CD44 expression in human ESF and SF. Note that the predicted CD44 expression is close to the actually measured expression in cultured human cells.

### Synthetic Interpretation of CD44 data

Comparative data of CD44 expression in SF and ESF suggests a concerted increase in the human lineage leading to similar levels of RNA expression in both human cell types (Figure 1b). This pattern is unexpected, as the variation of CD44 expression between cell types is in general quite independent. The correlation of phylogenetic independent contrast shows a modest correlation of 0.673, but this value is driven by the expression in human cells. Eliminating the contrast including the human cells reduces the correlation to 0.143, hence in all other species but human the evolution of CD44 expression is independent between cell types. CD44 expression in humans is not only more similar between cell types but also higher than in the other species investigated. This raises the question how and why this coordinated increase in CD44 expression evolved.

Compared to the most recent common ancestor of boreoeutherians the expression of CD44 increased along the human lineage by about 3x in both cell types (ESF 3.29x, SF 2.86X, average 3.07x). In the bovine lineage CD44 expression in ESF remains near constant (1.24x) and decreases in SF (0.26x). Reporter gene expression driven by the 3kb proximal CRE in cognate ESF reproduces the species difference: 2.95x difference of reporter gene expression between BtaCRE in BtaESF compared to HsaCRE in HsaESF, which is close to the 2.66x difference between native CD44 expression in BtaESF and HsaESF, as well as 3.29x inferred evolutionary change in the human ESF lineage (Figure 4A). Furthermore the bovine CRE drives reporter gene expression at the same level in both human celltypes (2.22x in HsaESF and 2.19x in HsaSF relative to TPB) (Table 3, line 1). These data suggest that the cow CRE, BtaCRE, is a reasonable proxy for the cis-regulatory activity in the ancestral genome before the concerted evolution of high CD44 expression in the human lineage.

**Table 3:**
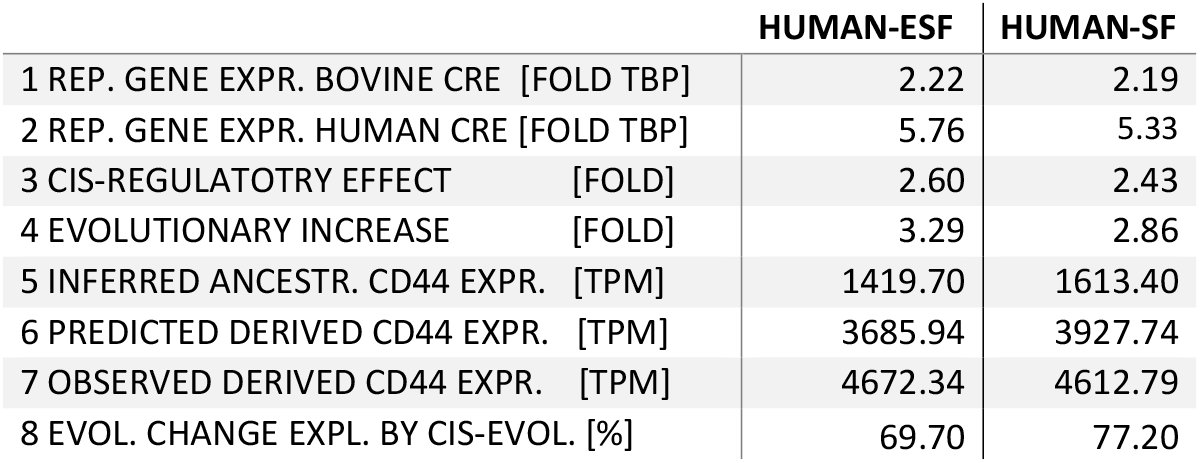
Comparison of experimental reporter gene results in human cell types (endometrial stromal fibroblasts ESF, skin fibroblasts SF) with evolutionary change in the human lineage of CD44 expression. Lines 1 and 2 are reporter gene expression values driven by bovine and human cis-regulatory elements [fold difference relative to TBP]; Line 3 is the inferred cis-regulatory effect (dividing value in line 2 by value in line 1); Line 4 is the fold increase in CD44 expression by dividing the observed DC44 expression (line 7) by the inferred ancestral CD44 expression (line 5); Line 5 is the inferred ancestral CD44 expression in the boreoeutherian ancestor (Figure 1b); Line 6 gives the predicted evolutionarily derived CD44 expression by multiplying the cis effect (line 3) with the inferred ancestral gene expression (line 5); Line 7 gives the CD44 gene expression in cultured human ESF and SF; Line 8 is the % of evolutionary gene expression explained by the cis-regulatory effect comparing the predicted derived CD44 expression (line 6) minus ancestral expression (line 4) with the difference between observed derived gene expression (line 7) minus ancestral gene expression (line 5).

|  | HUMAN-ESF | HUMAN-SF |
| --- | --- | --- |
| 1 REP. GENE EXPR. BOVINE CRE [FOLD TBP] | 2.22 | 2.19 |
| 2 REP. GENE EXPR. HUMAN CRE [FOLD TBP] | 5.76 | 5.33 |
| 3 CIS-REGULATORY EFFECT [FOLD] | 2.60 | 2.43 |
| 4 EVOLUTIONARY INCREASE [FOLD] | 3.29 | 2.86 |
| 5 INFERRED ANCESTR. CD44 EXPR. [TPM] | 1419.70 | 1613.40 |
| 6 PREDICTED DERIVED CD44 EXPR. [TPM] | 3685.94 | 3927.74 |
| 7 OBSERVED DERIVED CD44 EXPR. [TPM] | 4672.34 | 4612.79 |
| 8 EVOL. CHANGE EXPL. BY CIS-EVOL. [%] | 69.70 | 77.20 |

The effect of replacing the bovine reporter construct by its human counterpart leads to a similar cis-regulatory effect in both human cell types (2.60X in HsaESF and 2.43x in HsaSF, average 2.51x) (Table 3, line 3). We can use these measured cis-regulatory effects to estimate how much of the evolutionary change in the human lineage can be explained by the differences between the human and bovine CRE. Such a predicted change can be obtained by multiplying, for each cell type, the inferred ancestral gene expression level (Figure 1b) with the cis-regulatory effect measured in the reporter gene experiment (Table 3, line 3) and compare it with the measured CD44 expression in human cells (Figure 5B). This exercise suggests that 70 to 80%, depending on cell type, of the evolutionary change in the human lineage can be explained by the differences between bovine and human proximal CRE (Table 3, line 8). This inference implies that the majority of the changes in the human lineage are due to cis-regulatory effects located within 3kb upstream of the TSS of CD44.

It is difficult to formally assess whether additional cis-regulatory changes, not captured by the reporter construct, or trans-regulatory changes are also contributing to the human derived gene expression level not explained by the calculations in Table 3. The difficulty arises from modeling the error propagation when combining experimental results and inferences from comparative data. However, given that the reconstruction of the ancestral states used in these calculations is about +/- 20% of the estimated ancestral value, the formal confidence intervals of the predicted evolutionary outcome and the measured CD44 expression are overlapping, suggesting that this data does not support a strong contribution of other factors than those captured in the promoters captured in the reporter gene constructs.

What is the evolutionary scenario that is consistent with or suggested by these data? From the large fraction of evolutionary change explained by cis-regulatory differences it is tempting to conclude that the coordinated evolution of CD44 expression in ESF and SF is explained by pleiotropy, i.e. the assumption that mutations in the proximal CRE of CD44 always cause highly correlated gene expression changes in both cell types. However, this hypothesis is contradicted by the low correlation of CD44 expression in ESF and SF between species (Figure 1b) and the fact that one human lineage specific cis-regulatory change only affects gene expression in SF but not ESF. That is the acquisition of transcription factor binding sites for CEBPB in the human lineage (Table 2). However, KD of CEBPB only affects CD44 expression in SF but not in ESF (Figure 5A), which explains 50% of the evolutionary change in SF (Suppl. Figure 9). Given these observations, the more likely scenario is that the human lineage experienced selection in favor of increased CD44 expression in both cell types, i.e. correlative selection. The functional significance of higher CD44 expression for both cell types, and may be other stromal cell types, remains unclear. One speculative possibility is that the primate lineage experienced selection in favor of anti-fibrotic phenotype in all stromal cells, as CD44 interacts with the TGFb1 receptor inducing an anti-fibrotic phenotype (***). Increased CD44 expression could affect wound healing in the skin as well as non-scarring regeneration of the endometrium after menstruation or birth. Menstruation, as increased CD44 expression, evolved before the most recent common ancestor of apes and old world monkeys. It is intriguing to note that the only rodent species known to menstruate, the spiny mouse (*Acomys cartari*) is also known for extreme levels of skin regeneration. The increased vulnerability to cancer metastasis that also is associated with higher stromal CD44 expression would affect mostly later stages in life and thus have limited impact on fitness, as predicted by life history theory (Williams, ***).

## Conclusions

We conclude that the primate lineage evolved an increased expression of *CD44* in skin fibroblasts in concert with *CD44* expression in endometrial fibroblasts, while the expression in non-primates remains about three to four-fold lower and does not display distinct evolutionary trends or correlated, between cell types, expression change in the taxon sample investigated here. Higher *CD44* expression is causing higher stromal invasibility of fibroblast populations by cancer cells (Kshitiz, Afzal et al. 2019), and may thus, in part, explain the high vulnerability of humans to malignancies of the skin compared to that of bovines and horses (D’Souza and Wagner 2014). The dominant isoforms are the same in human and cattle, but there is a difference in iso-form composition with humans expressing a short transcript of only eight exons, HsaCD44-205, not described nor found by us in the cow. Transcription of *CD44* in both cell types and species is initiated from homologous promoters. Thus the concerted increase of gene expression in the human lineage can be due to mutations in this shared promoter sequence. Reporter gene assays and ancestral state reconstruction show that the differences between ancestral expression level and the derived one in the human lineage are predominantly caused by cis-regulatory changes

Overall the results supports a model where in the primate lineage the invasibility of stromal tissue has dramatically increased, most likely after the most recent common ancestor of Euarchontoglires (roughly primates and rodents) and before the ancestor of Cartarrhini, i.e. the clade of old world monkeys and apes, which includes humans. It is unclear what evolutionary processes have driven this evolutionary change in *CD44* expression. A notable association is that in the same (relatively wide) phylogenetic window evolved spontaneous decidualization in primates (Emera, Romero et al. 2012). Spontaneous decidualization means that the endometrium differentiates into decidua after ovulation even in the absence of an embryo as is the case in women, for instance, but only a small fraction of mammalian species have this characteristic (Critchley, Babayev et al. 2020). Menstruation is a consequence of spontaneous decidualization if no fertilization is happening. This characteristic can be interpreted as a maternal defense against highly invasive placentation (Finn 1998), i.e. compensating for the higher invasibility of the uterine stroma that evolved in the primate lineage.

We identified cis-regulatory changes in the *CD44* proximal promoter that explain almost 50% of the increase of CD44 expression in skin fibroblasts. Specifically, we found three binding sites for CEBPB at the human locus and almost none in the other species, and the knockdown of CEBPB reduces CD44 expression by about 30%.

The relatively large and cell type-specific contributions of trans-regulatory factors in SF suggests that it may be possible to manipulate *CD44* expression in the tumor-associated stroma cells with minimal side-effects on other cells and tissues. Such an intervention would be clinically desirable to prevent cancer cell dissemination from the site of the primary tumor.

## Materials and Methods

### Cell sourcing

#### Human Endometrial Stromal Cells and Skin Fibroblasts

Human ESFs were obtained from the Gil Mor group. Human SFs (BJ5ta) were purchased from ATCC (CRL-4001).

##### Skin Fibroblasts

Cow (*Bos taurus*), dog (*Canis lupus*), guinea pig (*Cavia porcellus*), horse (*Equus caballus*), cat (*Felis catus*), monkey (*Macaca mulatta*), opossum (*Monodelphis domestica*), rabbit (*Oryctolagus cuniculus*), sheep (*Ovis aries*), rat (*Rattus norvegicus), and* pig (*Sus scrofa*) SFs were obtained from fresh skin tissue. A small piece of skin was collected, hair removed and the sample was washed in PBS buffer and cut into strips approximately 1.0 cm2. Dermis was separated from epidermis by enzymatic digestion (30 min in 0.25% Trypsin buffer at 37 °C, followed by dissociation buffer (1 mg ml–1 collagenase, 1 mg ml–1 Dispase, 400 μg ml–1 DNase I) for 45 min at 37 °C). Epidermis was removed and 2 mm pieces were cut from the dermis and transferred to a 12-well plate and covered with media. Fibroblasts emerged from the explants and grew to confluency in growth media. Extra tissue was removed.

##### Endometrial Stromal Fibroblasts

Cow (*Bos taurus*), dog (*Canis lupis*), guinea pig (*Cavia porcellus*), horse (*Equus caballus*), cat (*Felis catus*), monkey (*Mucaca mulatta*), opossum (*Monodelphis domestica*), rabbit (*Oryctolagus cuniculus*), sheep (*Ovis aries*), rat (*Rattus norvegicus), and* pig (*Sus scrofa*) ESFs were obtained as follows. Uterine tissues were collected from each species and primary ESFs were obtained by enzymatic digestion. Uterus fragments, 2–3 mm in size, were created using a scalpel and digested with 0.25% Trypsin–EDTA for 35 min at 37 °C, followed by dissociation buffer (1 mg ml–1 collagenase, 1 mg ml–1 Dispase, 400 μg ml–1 DNase I) for 45 min at 37 °C. Cell clumps were homogenized by passage through a 22-gauge syringe followed by passage through a 40-μm nylon mesh filter to remove remaining clumps. For all species except opossum, a single-cell suspension was obtained from the lysate, transferred to fresh growth medium and cultured in T25 flasks. To facilitate enrichment of fibroblasts versus epithelial cells, media were exchanged in each well after 15 min to remove floating cells that had not yet attached while stromal fibroblasts had attached. Cells were grown to confluency and subpassaged by scraping the cells off the surface to be split into two T25 flasks. For opossum, the single cell suspension was layered onto a percoll density gradient for further separation. Immunohistochemistry was used to test for abundance of vimentin (Santa Cruz, sc-6260) and cytokeratin (Abcam, ab9377) to validate fibroblast subtype in the isolated cells.

### Cell culture

ESFs were grown in phenol-red free DMEM/F12 with high glucose (25 mM), supplemented with 10% charcoal-stripped calf serum (Gibco) and 1% antibiotic/antimycotic (Gibco). BJ5ta (ATCC) cells were cultured in 80% DMEM and 20% MEM supplemented with 10% FBS, 1% antibiotic/antimycotic and 0.01 mg ml–1 hygromycin. SFs were cultured in DMEM with high glucose supplemented with 10% FBS.

### RNA isolation and sequencing

RNA was isolated using RNeasy micro kit (QIAGEN) and resuspended in 15 μl of water. The Yale Center for Genome Analysis ran samples on the Agilent Bioanalyzer 2100 to determine RNA quality, prepared mRNA libraries and sequenced on Illumina HiSeq2500 to generate 30–40 million reads per sample (Single-end 75 base pair reads).

### Transcript-based abundances using RNAseq data

RNAseq data obtained was quantified using the transcript-based quantification approach as given in the program ‘kallisto’ (Bray, Pimentel et al. 2016). Here reads are aligned to a reference transcriptome using a fast hashing of k-mers together with a directed de Bruijn graph of the transcriptome. This rapid quantification technique produces transcript-wise abundances which are then normalized and mapped to individual genes and ultimately reported in terms of TPM (Li, Sun et al. 2009, Wagner, Kin et al. 2012). The Ensembl release 99 (Cunningham, Achuthan et al. 2019) gene annotation model was used and raw sequence reads (single end 75 bp) for ESFs and SFs from human (*Homo sapiens*), cow (*Bos taurus*), dog (*Canis lupus*), cat (*Felis catus*), guinea pig (*Cavia porcellus*), horse (*Equus caballus*), monkey (*Macaca mulatta*), opossum (*Monodelphis domestica*), rabbit (*Oryctolagus cuniculus*), sheep (*Ovis aries*), rat (*Mus musculus*), pig (*Sus scrofa*) were aligned to GRCh38.p13, ARS-UCD1.2, CanFam3.1, Felis_catus_9.0, Cavpor3.0, EquCab3.0, Mmul_10, MonDom5 (Release 97), OryCun2.0, Oar_v3.1, Rnor_6.0 and Sscrofa11.1 reference transcriptome assemblies. In order to facilitate gene expression across species, a one-to-one ortholog dataset consisting of 8639 species was formulated (comprising of human, rat, rabbit, horse, guinea pig, cat, sheep, cow, horse, dog and opossum) such that the sum of TPMs across these 8639 genes for each species totals to 10^6^. Additionally, the extended SF dataset comprising of pig and monkey was formulated and contained a total of 7743 one-to-one orthologs. Ancestral character estimation was performed on sqrt(TPM) using the Residual Maximum Likelihood method (REML) available in the ‘ace’ module of APE (Paradis, Claude et al. 2004) in R statistical package.

### Isoform abundances using RNAseq data

The reads obtained as a result of RNA sequencing were quantified by adopting a transcript-based quantification approach. The RNAseq quantification program called Kallisto (Bray, Pimentel et al. 2016) was utilized to this end wherein the reads were aligned to an indexed reference transcriptome using a fast hashing of k-mers together with a directed de Bruijn graph of the transcriptome (Compeau, Pevzner et al. 2011). A list of transcripts that are compatible with a particular read are generated. This rapid quantification technique produces transcript-wise abundances and these were reported in terms of TPM (Li, Sun et al. 2009, Wagner, Kin et al. 2012). In order to identify splice variants from RNAseq data, we utilized transcript-based abundances obtained from Kallisto. A comprehensive literature survey on exon architecture of CD44 was followed by mapping the Kallisto-obtained abundances to reported CD44 isoforms in order to verify which splice variants are expressed.

### Protein abundances

We used a proteomic method called data-independent acquisition mass spectrometry (DIA-MS) (Aebersold and Mann 2016) to quantify the ratio of CD44 proteins from each sample to the total proteomes. The implementation of DIA-MS was identical to published (Mehnert, Li et al. 2019). Briefly, the human and cow skin fibroblast cell samples were washed, harvested, and snap-frozen by liquid nitrogen. The protein extraction was performed by adding 10 M urea containing complete protease inhibitor cocktail (Roche) and Halt™ Phosphatase Inhibitor (Thermo) and digested. About 1.5 micrograms of peptides from each sample were used for DIAMS measurement on an Orbitrap Fusion Lumos Tribrid mass spectrometer (Thermo Scientific) platform coupled to a nanoelectrospray ion source, as described previously (Mehnert, Li et al. 2019). DIA-MS data analyses were performed using Spectronaut v13 (Bruderer, Bernhardt et al. 2017), by searching against the UniProt proteome databases of Homo sapiens (human) and Bos Taurus (cow) separately for samples of different speicies. Both peptide and protein FDR cutoff (Qvalue) were controlled at 1%, and the label-free protein quantification was performed using the default settings in Spectronaut.

### CD44 Methods

#### 5’-RACE

RNA was isolated by RNeasy mini kit (Qiagen) from BJ5ta cells, bovine dermal fibroblasts, human endometrial stromal fibroblasts, and bovine endometrial cells. RNA was purified via phenol-chloroform extraction. The mRNA was enriched using the MagJET mRNA Enrichment Kit (Thermo Scientific). Then, the 5’-RACE procedure was performed using the FirstChoice^®^ RLM-RACE Kit (Invitrogen™). cDNA containing the 5’UTR of CD44 from each cell type was amplified using a nested PCR strategy with the primers:

Arbitrary outer, forward adapter (both human and bovine CD44):
GCTGATGGCGATGAATGAACACTG
Arbitrary inner, forward adapter (both human and bovine CD44): ACTGCGTTTGCTGGCTTTGATG
Human CD44 outer, reverse: GGAGGTGTTGGATGTGAGGATGTA
Human CD44 inner, reverse: CATTGTGGGCAAGGTGCTATTG
Bovine CD44 outer, reverse: GGAGGTGTTGGATGTGAGGATGTA
Bovine CD44 inner, reverse: ATGGTGGGCAGCGTGCTATTA

PCR products were run on 2% SDS-polyacrylamide gels, gel-extracted, sequenced, and aligned to the human or bovine genome to elucidate the 5’-UTR length and, ultimately, the transcription start site of CD44 in each cell type.

### Analysis of cis-regulatory elements

We searched for putative cis-regulatory elements that may affect CD44 gene expression n the different species. Position-specific frequency matrix motifs of known eukaryotic transcription factor binding sites were downloaded from the JASPAR database (Khan, Fornes et al. 2018). The genome sequences of the species were obtained from the Ensembl database: https://doi.org/10.1093/nar/gkz966. We considered genomic regions 5kb upstream to 1kb downstream of the translation start site of each gene as the promoter regions to be analyzed. Alignment to the binding site motifs within the CD44 promoter region in each species was calculated using the FIMO package using default parameters (Grant, Bailey et al. 2011). We considered valid matches where the alignment was reported to be statistically significant with a p-value < 10^−4^. CEBPB binding sites were then selected for further analysis.

### Plasmids and Constructs

The promoter fragment (−2,679−+406) of The human CD44 was PCR amplified from chromosomal DNA of Human ESF cells with primers introducing XhoI (5’-GCCG<u>CTCGA</u>GAGGTTCCATGAAACACAGTAAGA -3’) or HindIII(5’-CCC<u>AAGCTT</u>GCGAAAGGAGCTGGAGGAT -3’) restriction sites. The cow CD44 promoter fragment (−2,886−+42) was PCR amplified from chromosomal DNA of Cow ESF cells with primers introducing XhoI (5’-CCG<u>CTCGAG</u>CTGCTAAGTCGCTTCAGTCAT -3’) or HindIII(5’-AGCCC<u>AAGCTT</u>GGAAGTTGGGTGCAGlllll -3’) restriction sites. The resulting fragment was cloned into a Promoterless NanoLuc^®^ Genetic Reporter Vectors pNL2.1[Nluc/Hygro] (Promega), and sequence was verified.

### Transfection and Luciferase Assays

For CD44 promoter analysis, cells in 24-well plates were co-transfected with 200 ng human or cow CD44-promoter-pNL2.1 or empty vector pNL2.1, and 40 ng pGL4.13 with Lipofectamine 3000 (Invitrogen) according to the manufacturer. cells were lysed 24 hours after transfection for luciferase assay with the Nano-Glo Dual-Luciferase^®^ Reporter Assay System (Promega). CD44 promoter activity (NanoLucR luciferase) was normalized with firefly luciferase activity.

### Real-Time PCR

Cells in 6-well plates were transfected with 1 ug human or cow CD44-promoter-pNL2.1 or empty vector pNL2.1 using Lipofectamine 3000 (Invitrogen) as manufacturer’s instructions. Total RNAs from cells were isolated using RNeasy Plus Micro Kit (QIAGEN) according to the manufacturer’s protocol. After digestion by RNase-Free DNase Set (QIGEN), RNAs were reverse transcribed according to iScript cDNA Synthesis Kit (Thermo Fisher Scientific). Real-time polymerase chain reaction (PCR) was performed using an Applied Biosystems™ Fast SYBR™ Green Master Mix (Thermo Fisher Scientific). Primer sequences used for real-time PCR are Nluc-1F:CAGGGAGGTGTGTCCAGTTT, Nluc-1R:TCGATCTTCAGCCCATTTTC for evaluation of CD44-promoter-driven Luciferase activity; Hygr-1F:GAGCCTTCAGCTTCGATGTC, Hygr-1R:CGGTACACGTAGCGGTCTTT for evaluation of the expression of hygromycin driven by constitutive promoter. Hygromycin expression was used for normalization. The 2−ΔΔCt method was used for data analysis.

For detecting endogenous cow and human CD44 expression, the primer sequences used for real-time PCR were listed as follows:

Bt-CD44-Forwad: TACAGCATCTTCCACACGCA,
Bt-CD44-Reverse: GCCGTAGTCTCTGGTATCCG
Bt-TBP-2F:GCACAGGAGCCAAGAGTGAA,
Bt-TBP-2R:TTCACATCACAGCTCCCCAC
Hs-CD44-1F: GATGGAGAAAGCTCTGAGCATC
Hs-CD44-1R: TTGCTGCACAGATGGAGTTG
Hs-TBP-1F: GGAGAGTTCTGGGATTGTAC
Hs-TBP-1R:CTTATCCTCATGATTACCGCAG
TBP was used for normalization.

### Cell Culture, RNA-interference, & Quantitative PCR

Human Skin Fibroblasts (BJ-5ta, ATCC CRL-4001) were cultured in a 4:1 mixture of 4 parts Dulbecco’s Modified Eagle’s Medium containing 4mM L-glutamine, 4.5g/L glucose and 1.5g/L sodium bicarbonate and 1 part Medium 199 supplemented with 0.01mg/mL hygromycin B, and 10% fetal bovine serum.

Human Endometrial Stromal Fibroblasts (T-HESCs, ATCC CRL-4003) were cultured in a 1:1 mixture of Dulbecco’s Modified Eagle’s medium and Ham’s F-12 medium with 3.1g/L glucose and 1mM sodium pyruvate without phenol red (D2906, Sigma) supplemented with 1.5g/L sodium bicarbonate, 1% ITS+ Premix (354352, BD), 1% ABAM, and 10% Charcoal stripped fetal bovine serum (100-119, Gemini).

12-well plates were grown to 70% confluency and transfected with 25nmol final concentration of siRNAs targeting CEBPb (s2891 and s2892 Themo Fisher). In preparation for transfection, siRNAs in OptiMem I Reduced Serum Media (31985, Thermo Fisher) were mixed with an equal volume of OptiMem containing Lipofectamine RNAiMax (13778, Thermo Fisher), incubated at room temperature for 20 min, and added dropwise to cells in 1 ml growth media. Final concentration of siRNAs was 25 nM. Control wells were prepared without any siRNA added.

Knockdown of CEBPb and expression of CD44 were confirmed by qPCR. Media was removed, cells were washed in PBS followed by direct lysis with Buffer RLT Plus + beta-mercaptoethanol. RNA was extracted according to the manufacturer’s protocol (74034, RNeasy Plus Micro Kit, Qiagen). Reverse transcription of 1 μg of RNA was carried out with iScript cDNA Synthesis Kit (1708891, Bio-Rad) using an extended transcription step of three hours at 42°C. qPCR reactions were with Taqman Fast Universal PCR Master Mix (4366072, Applied Biosystems) in duplicate using 5 ng of cDNA for template each. Taqman probes for CEBPb (hs00270923_s1, Invitrogen) and CD44 (Hs01075864_m1, Invitrogen) were used to amplify each template. Fold change was calculated by finding the ddCt values relative to the expression of TATA Binding Protein.

## Appendix

### Defining interaction and independence

While the detection, by statistical means, of interactions between various experimental effects is a technique with a >100 year tradition, there still exists considerable confusion over how to define and detect interaction effects and how they can be interpreted. Since some of our results critically depend on the correct identification of such interaction effects, we want to briefly outline our approach, which was first developed in the context of fitness measurements (Wagner 2010) and is an extension of the theory of measurement and scale types (Nares 2002). This approach was generalized to the principle of “effect propagation” (Wagner 2015). In brief the proposal says, 1) effects need to be calculated in a way that respects the constraints of scale type, and 2) that the way effects are calculated determines the way interaction effects have to be defined. For instance, if experimental effects can be calculated as differences, then interaction effects have to be defined as deviations from additivity. In contrast, if the experimental effects are measured as fold changes (factors) then interaction effects are deviations from multiplicative composition. Formally the principle of effect propagation reads:

*Let V be a response variable in a factorial experiment, and X and Y some experimental manipulations or natural alterations (e.g. two mutations, or changing the cis-regulatory element, CRE, or the cell type in a reporter experiment). V(C) is the value in a reference or control experiment. Let m(X) and m(Y) be the direct effects of the manipulations X and Y, m(X)=f[V(C), V(X)] and m(Y)=f[V(C), V(Y)]. An effect measure can be introduced that represents the result of the experimental manipulations if there is a mathematical operation ∘ such that V(X) = V(C) ∘ m(X) and V(Y) = V(C) ∘ m(Y), and “∘ “ is invertible, associative and commutative. The combined effect of X and Y in the absence of interaction then is m(X&Y) = m(X) ∘ m(Y). The interaction effect then is the deviation between the measured combined effect of X and Y, m(X&Y), and the one calculated by m(X) ∘ m(Y), i.e. V(X&Y)≠V(C) ∘ (m(X) ∘ m(Y))*.

The formal proof can be found in Wagner (2015). Here the symbol “∘” stands for any mathematical operation that can be used to define an effect measure, i.e. addition/subtraction, multiplication/division or any other operation compatible with the scale type of V (for scale type constraints see for instance (Nares 2002)). For this study the implication is that since our qPCR results are expressed as fold differences between a target gene and a reference gene, the appropriate way to measure difference are fold changes. Interaction effects are then deviations from multiplicative combinations of the direct cis- and the trans-regulatory effects.

#### Multiplicative independence

Measuring the cis and the trans-effects we perform experiments where we measure the reporter gene expression, [RG], from a cis regulatory element of a species say a, CRE_a_, in a cell type CT_b_ from species b: CRE_a_ x CT_b_ and in all combinations of species a and b.

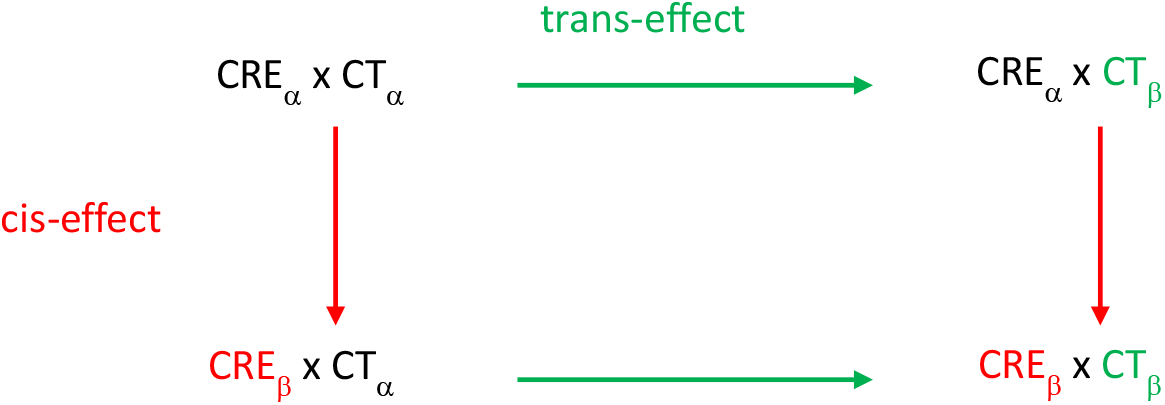

In terms of measured reporter gene expression this schema translates into:

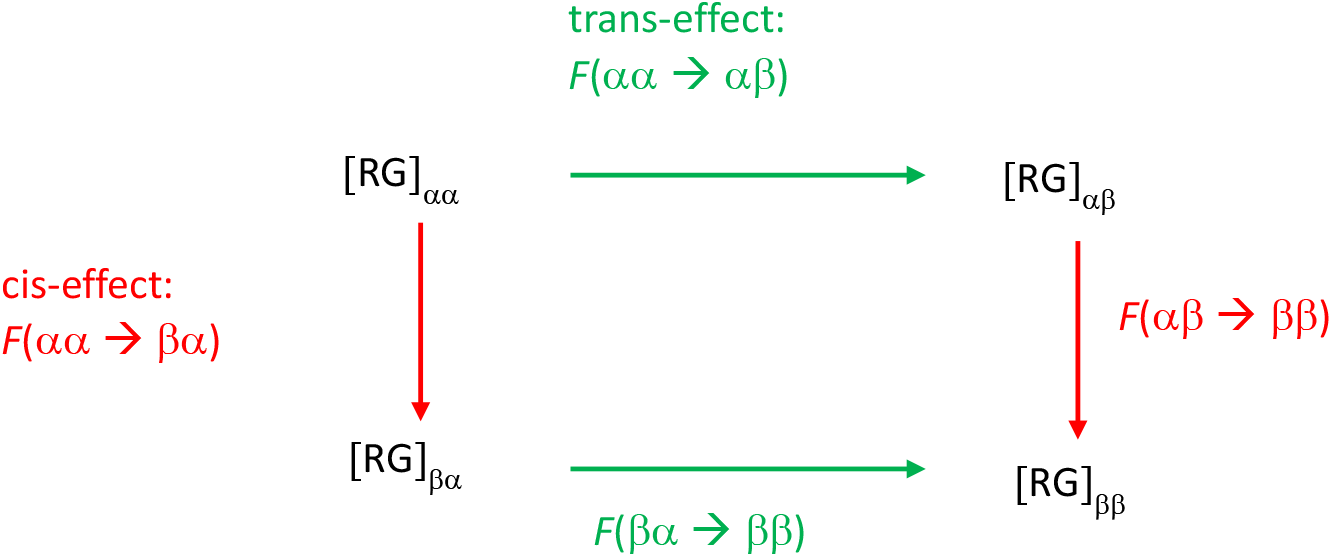

where the first index refers to the species of origin of the CRE and the second to the species of origin of the cell type in which the CRE is tested. The cis-effects and trans-effects are independent if they do not depend on the reference measurement. For instance, if the effect of substituting the CRE from human with that of the cow does not depend on whether this was done in a human or a cow cell.

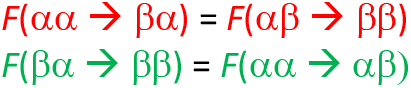

Similarly, the effect of changing the cell type from a human cell to a cow cell shall not depend on whether we test a human CRE or a cow CRE. Both of these equations imply the following relationship between the measured reporter gene expressions:

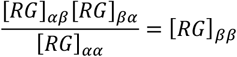

From this it is easy to see that if the cis and the trans-factors combine multiplicatively to determine the reporter gene expression level their effects are independent, i.e. there is no interaction. Below we summarize simple models of transcriptional regulation to determine whether we should expect independent cis- and trans-effects.

#### An “ideal gene theory”

For the sake of accessibility of our interpretation of the transfection data we provide the derivation of a simple kinetic model of gene regulation. The analysis her is inspired by the ideas in (Buchler, Gerland et al. 2003). We call this model the “ideal gene theory” to emphasize that this model is meant as a reference model, like the ideal gas model in physics, rather than a model of any real gene. Ideal reference models serve the purpose of a “foil” against which experimental data can be interpreted. Deviations of the data from the ideal model justify the inference that there has to be a relevant complicating factor. In order to allow this inference the ideal reference model has to be mathematically and conceptually explicit.

Let us consider a simple gene G that is the target of an activating transcription factor, TF. The transcription factor can bind to a cis-regulatory element with a rate *k’_1_* and dissociates from the binding site with rate *k_2_*. Let the probability of occupancy of the binding site be *P_1_* and the probability that the binding site is empty is *P_0_*. Then the rate of change of occupancy probability is described by the simple master equation:

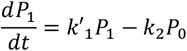

With *k’_1_= k_1_*[*TF*]. At equilibrium we have 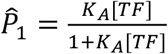 with *K_A_* is the association equilibrium constant 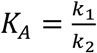. If *K_A_*[*TF*] = *q* ≪ 1, i.e., the binding site is far from saturation, the equilibrium probability of occupancy reduces approximately to 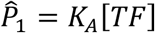.

Let us assume that indeed q<<1 and the transcription factor is an activator of transcription (rather than repressor). Let *k_tr_* be the rate of transcription when the CRE is occupied and *k_d_* the rate of mRNA degradation. Then the rate of change of the concentration of the mRNA corresponding to target gene sequence will be

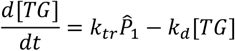

And in equilibrium we will have

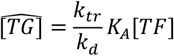

Or

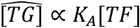

In interpreting our transfection experiments we interpret *K_A_* as the influence of the <u>cis-regulatory</u> element, and [*TF*] as representing the influence of the <u>trans-regulatory environment</u>, i.e. the cell. Therefore, cis- and the trans-environment multiplicatively determine the level of gene expression. As shown above, this model predicts that, under its conditions of validity, the cis- and trans-effects on reporter gene expression should show no interaction effect.

Now let us consider a model with two independently binding transcription factors, TF_1_ and TF_2_, with the equilibrium association constants *K_A1_* and *K_A2_*, *q_1_*=*K_A1_ [TF_1_]* and *q_2_*=*K_A2_ [TF_2_]*. Since each transcription factor acts independently the single occupancy probabilities are calculated in the same way as in the one TF case. Double occupancy probability P_3_ can easily be calculated based on independence of binding events as P_3_=P_1_ P_2_ and hence

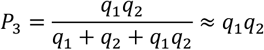

If *q_i_*<<1. In order to calculate the consequences for our reporter gene experiments we need to distinguish between two scenarios: an additive effect of transcription factor occupancy on transcription rate and a cooperative effect, where transcription only happens under double occupancy by both TFs. Let us assume two rate constants for each TF’s influence on reporter gene transcription, *k_tr1_* and *k_tr2_*. Assuming that both factors interact with the RNA polymerase complex independently, the equilibrium concentration of the reporter gene mRNA combining the effects of each factor separately, and their joint effect is

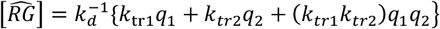

Since *q_i_*<<1 the transcription rate is dominated by the single occupancy events, which combine additively (one can neglect the last term), and thus the effect is expected to be additive.

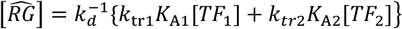

And cis- and trans-effects are not combining multiplicatively and will lead to interaction effects. Hence our result of independence of cis- and trans-effects refutes the idea that CD44 is controlled by multiple TFs acting independently and additively on the transcription rate.

In the case of cooperative transcriptional effects of the two transcription factors the equilibrium expression of the reporter gene is predicted to be:

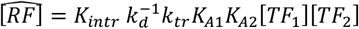

Here we introduced a new factor, K_int_, to account for the likely scenario of cooperative binding that is not independent (the binding of one factor increases the probability of the binding of the other factor). This assumption represents the likely scenario of why the term *K*_*intq*_1_*q*_2__ can be much greater than the terms *q_1_* and *q_2_*, even if *q_i_*<<1. The cis and trans effects are combining multiplicatively in this case. Hence the single TF and the multiple cooperative TFs models both predict no interaction between cis and trans-acting factors.

Finally, we consider another scenario, where we contrast the effects of different cell types, n and m, e.g. skin fibroblast (SF) or endometrial fibroblast (ESF), from different species, while keeping the CRE constant.

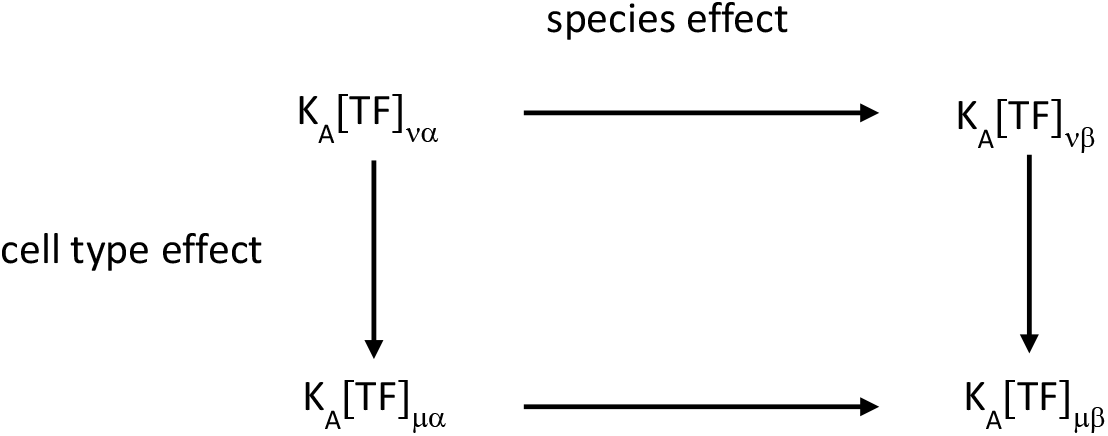

since we test the reporter gene expression with the same CRE, the cis-factor equilibrium association constant, *K_A_*, remains the same in all four measurements and can be dropped from the equations. The only differences are the cell type, e.g. SF and ESF, and the species, a and b, of origin of the cells. In this analysis, we compare four different trans-regulatory environments, for which our statistical analysis suggests a statistical interaction effect. The criterion for independence applied to this experimental setup then reads:

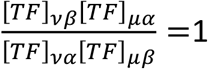

This condition can be interpreted in two ways. Rearranging the equation like this implies that the cell type effect in both species is the same.

The other re-arrangement leads us to this equation

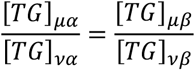

which implies that the species effect is the same for both cell types. This is perhaps the biologically more interesting statement, since a violation of this condition, i.e. a CT x species interaction effect, means that the trans-regulatory landscape in the two cell types evolved to some degree independently of each other. Hence the violation of this condition is prima facie evidence that the two cell types are in fact distinct cell types, i.e. evolve different trans-regulatory landscapes in different lineages, i.e. in the lineages of species a and b after the most recent common ancestor of these species.

## Supplementary Figure Captions

**Suppl. Figure 1:**
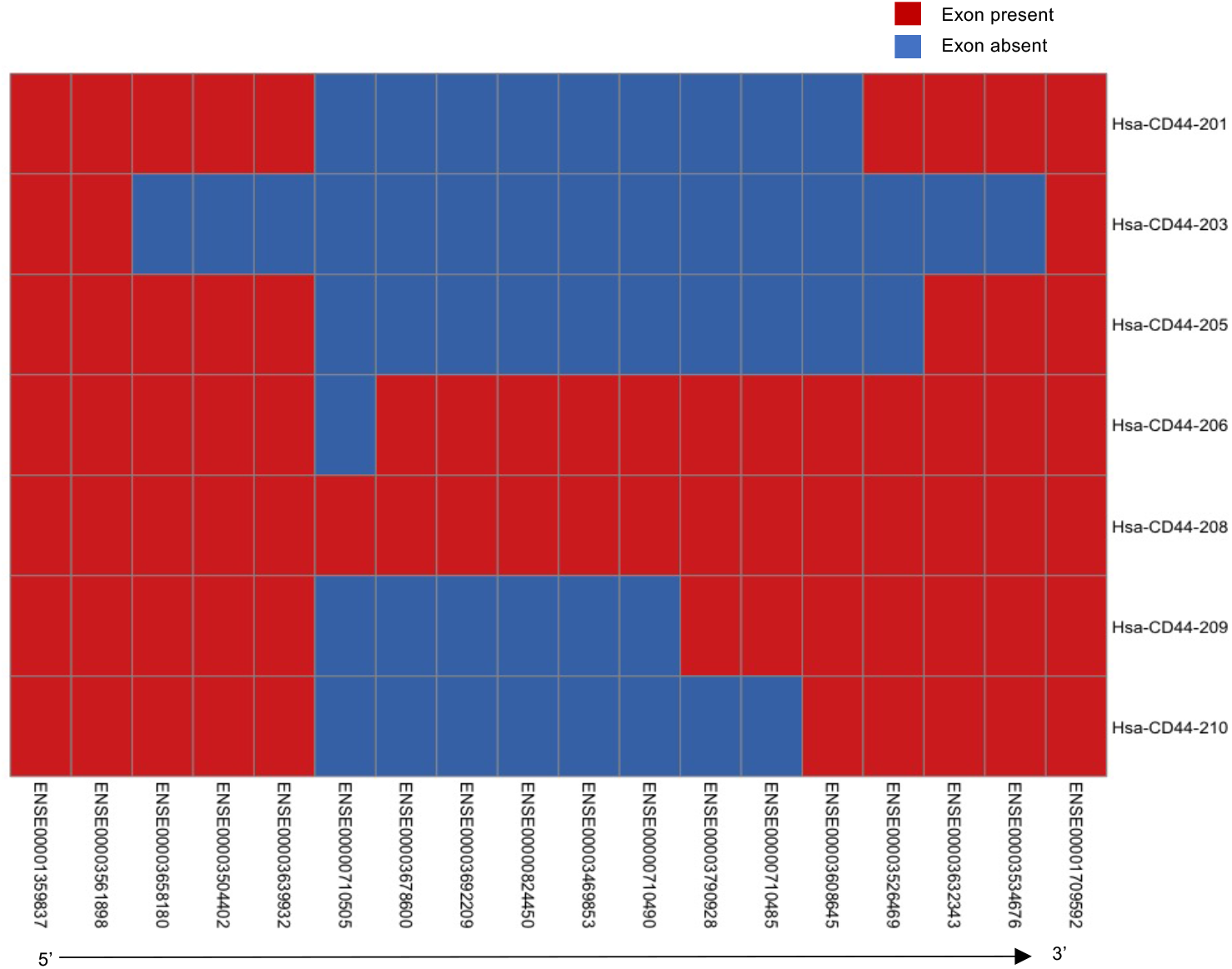
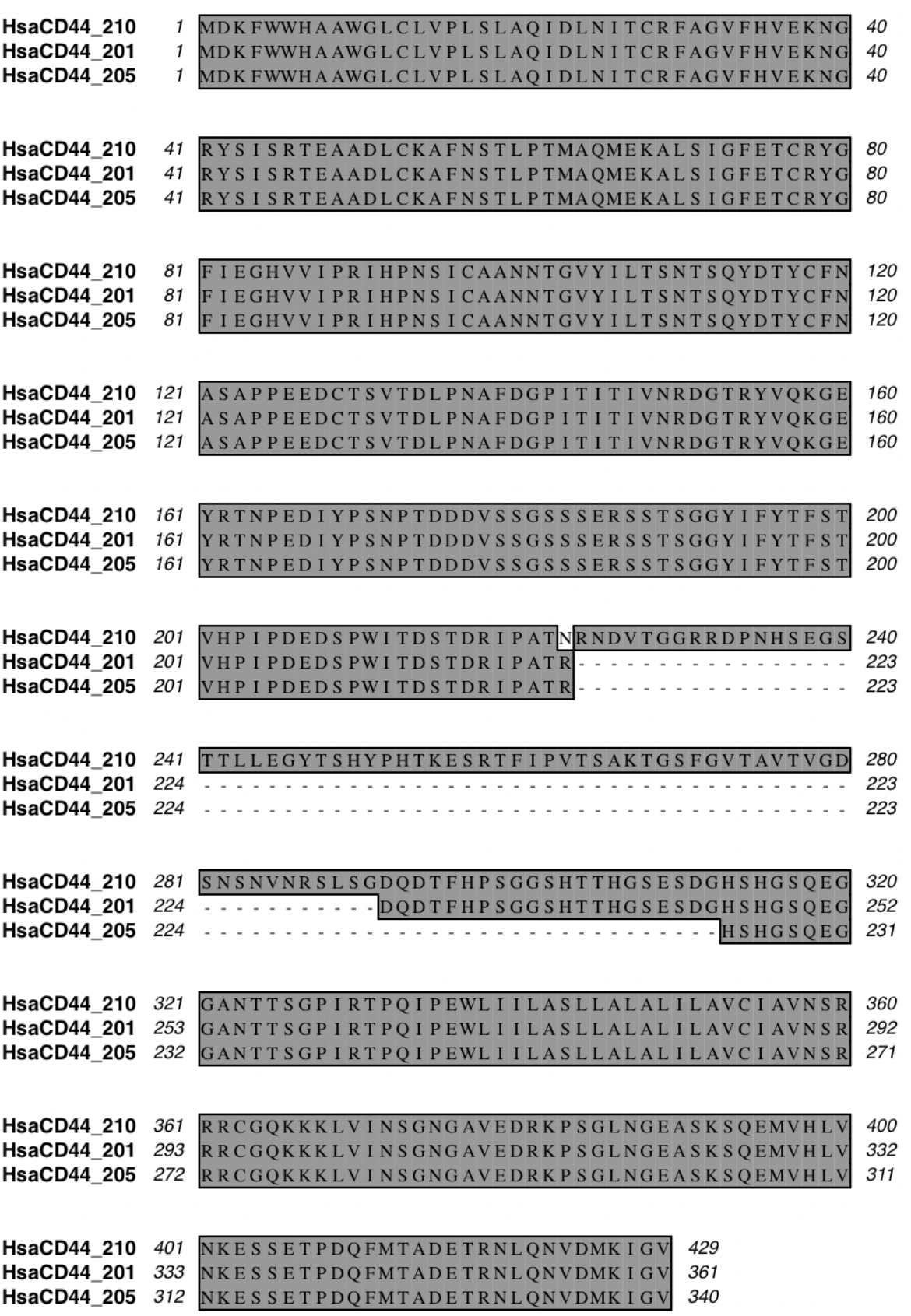
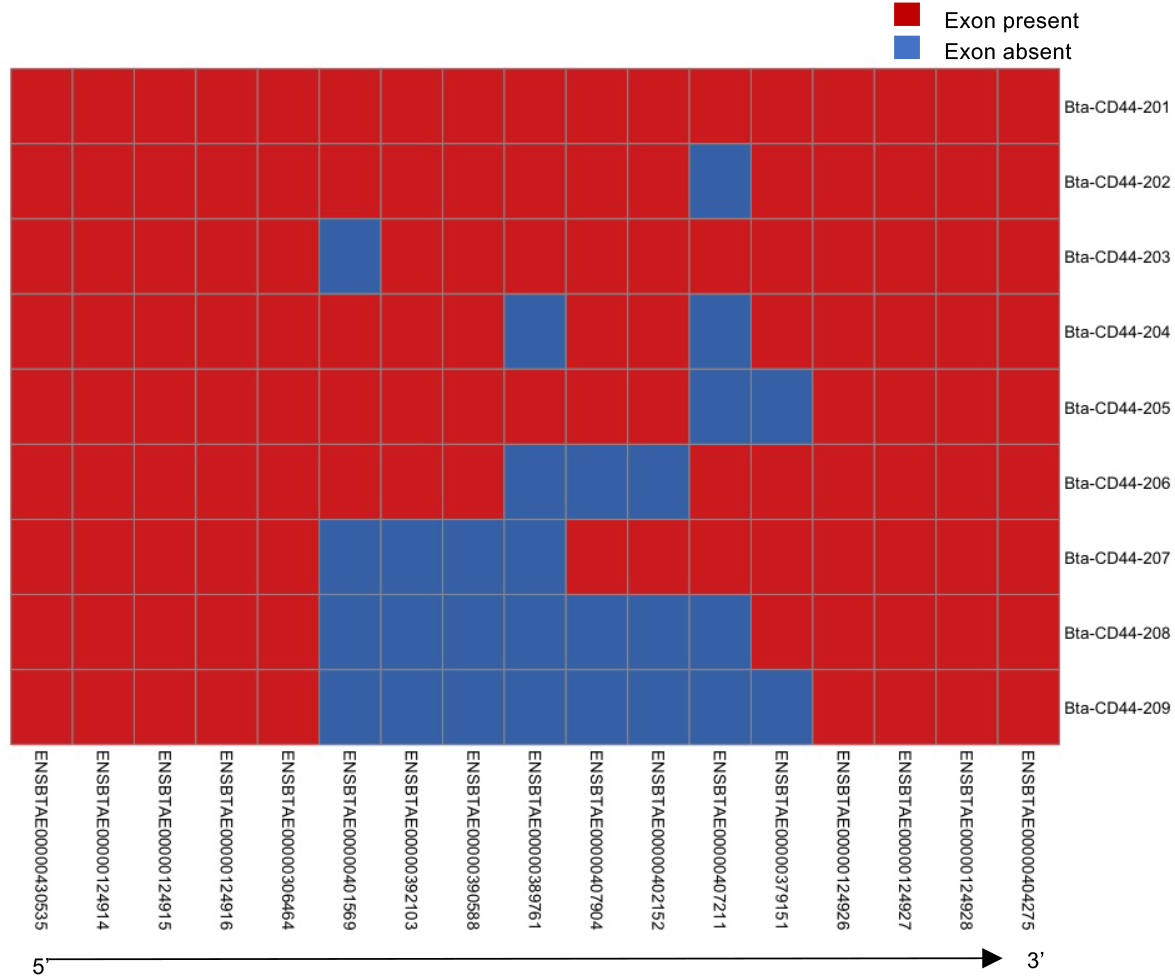
Exon composition of CD44 transcripts A) Exon composition of transcripts obtained from human SF and ESF RNA. B) Alignment of the computational translation of transcripts of the three dominant isoforms. The alignment shows that the different identities of the 3’ and 5’ most exons does not affect the amino acid sequence. C) Exon composition of transcripts identified from RNAseq data from cow mesenchymal cells.

**Suppl. Figure 2:**
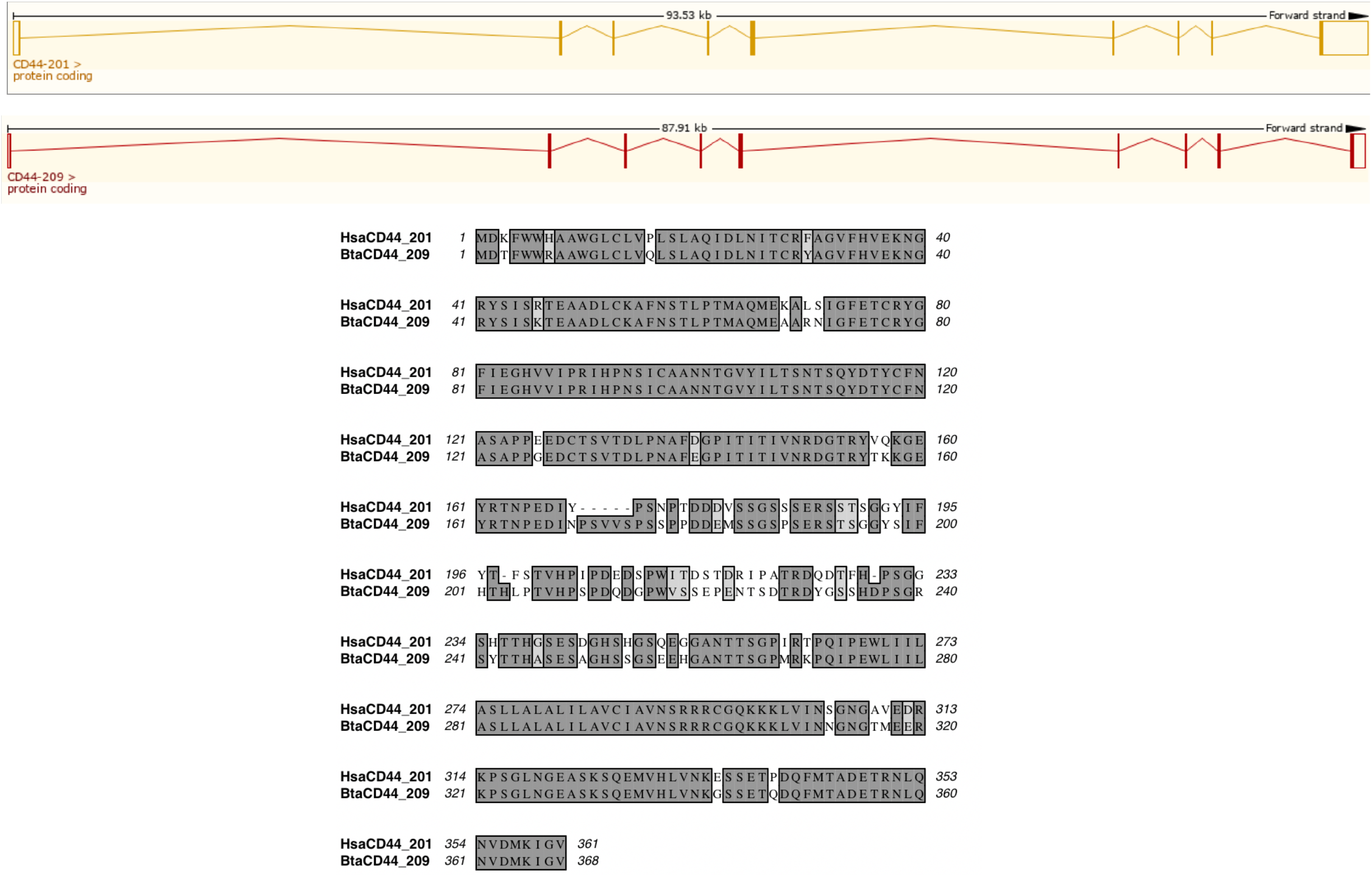
homology of HsaCD44-201 and BtaCD44-209. These transcripts consist or 9 exons and correspond to the human CD44s and the protein ID P16070-13 aka CD44R4.

**Suppl. Figure 3:**
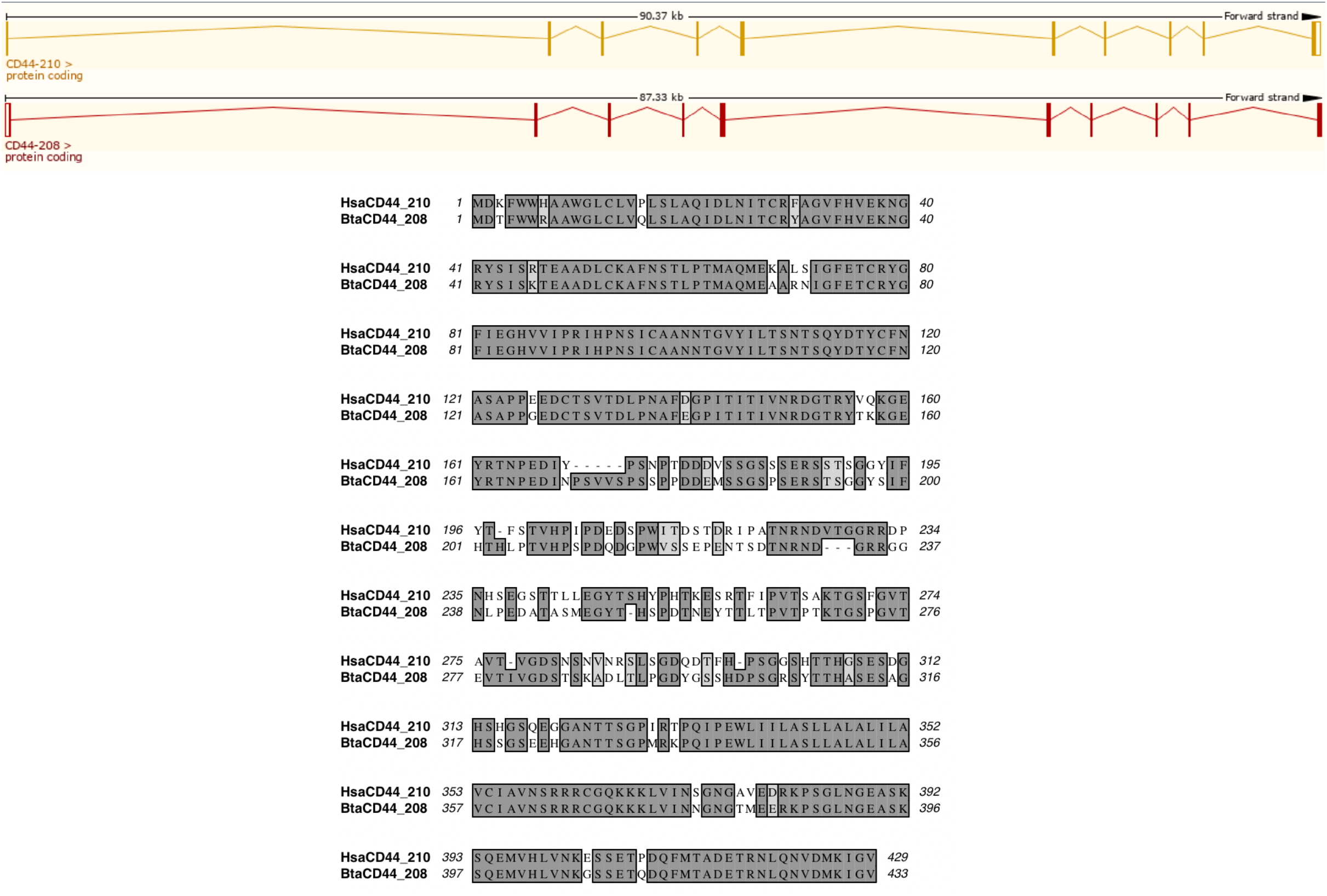
homology of HsaCD44-210 and BtaCD44-208. These transcripts consist of 10 exons including the variable exon 10, CD44v10, and has the protein ID P16070-11 aka CD44R2.

**Suppl. Figure 4:**
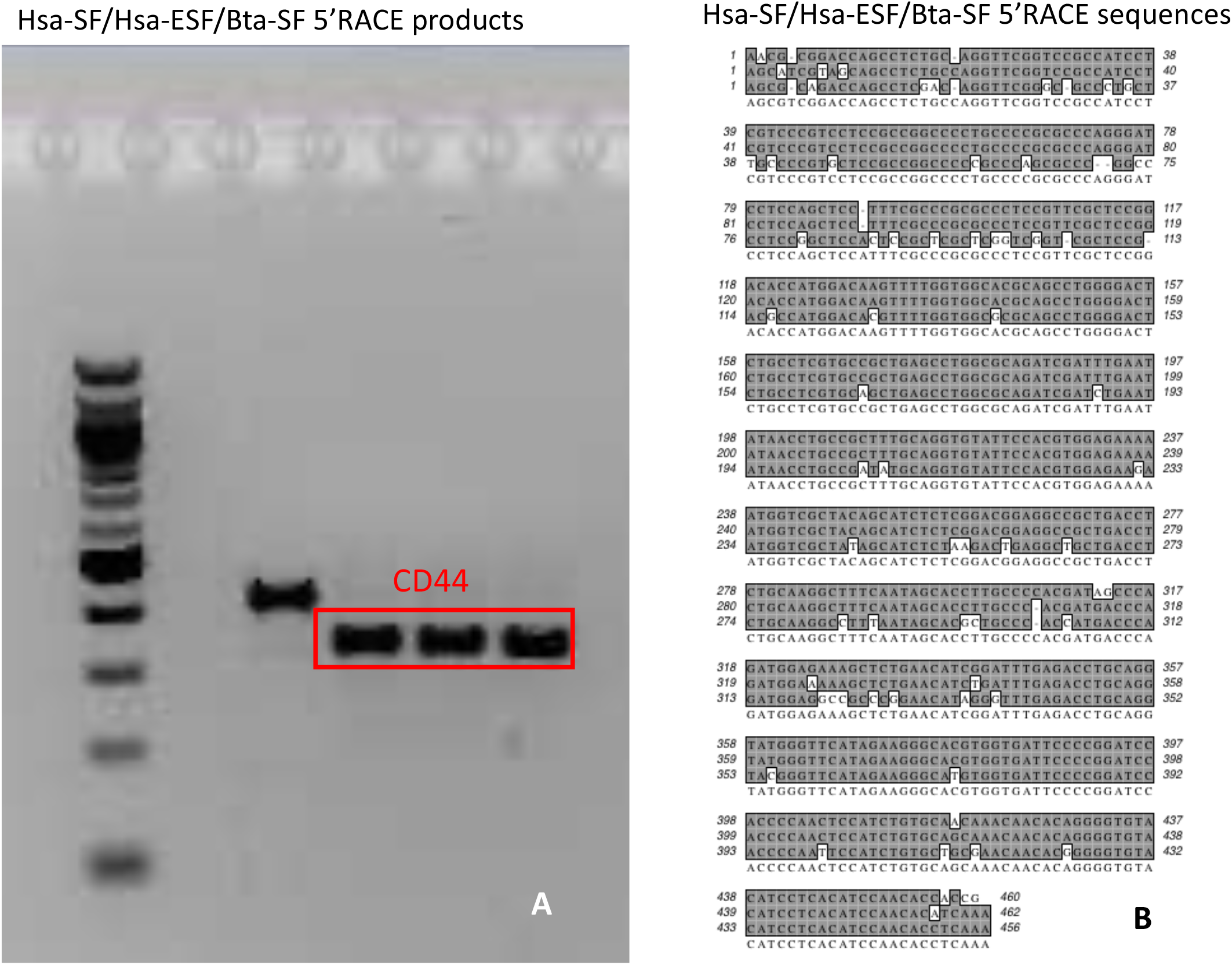
the 5’ RACE products of 5’ UTR of the human and cow CD44 from human ESF and SF as well as cow SF have the same length. A) gel images of 5’ RACE products B) alignment of the three RACE products.

**Suppl. Figure 5:**
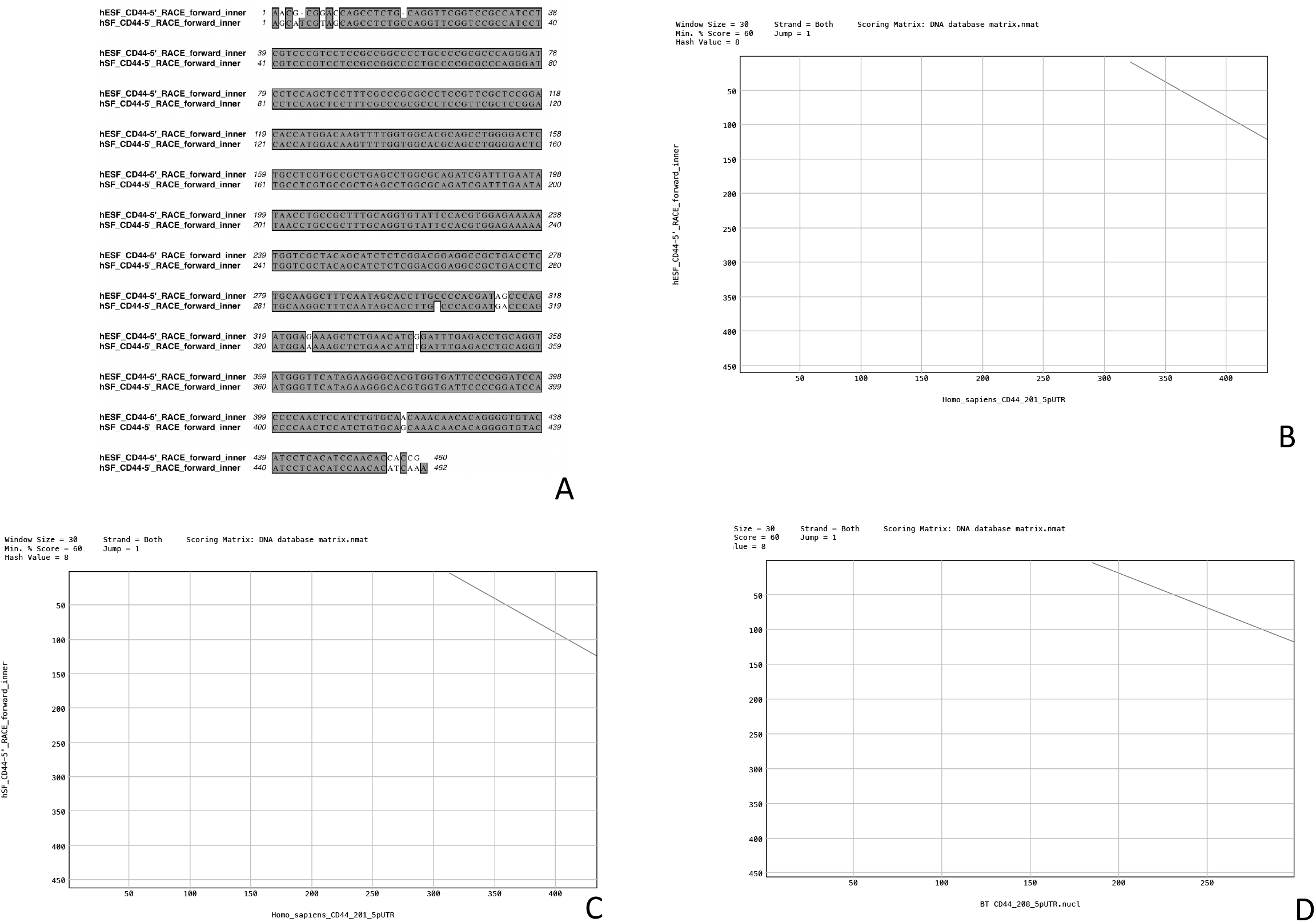
comparison of the human 5’ RACE products obtained in this study with the 5’ exons annotated in ENSEMLE. A) alignment of 5’ RACE products obtained from human SF and ESF showing that they are from the same dominant isoforms. B) dotplot of HsaCD44-201 5’ UTR with the RACE product from human ESF. The dotplot indicates that the longer 5’ UTR of HsaCD44-201 is overlapping with that of our RACE product, suggesting the use of an alternative promoter in the cells from which HsaCD44-201 was cloned. C) dotplot of HsaCD44-201 5’ UTR with the RACE product from human SF. D) dotplot of BtaCD44-208 5’ UTR with the RACE product from bovine SF.

**Suppl. Figure 6:**
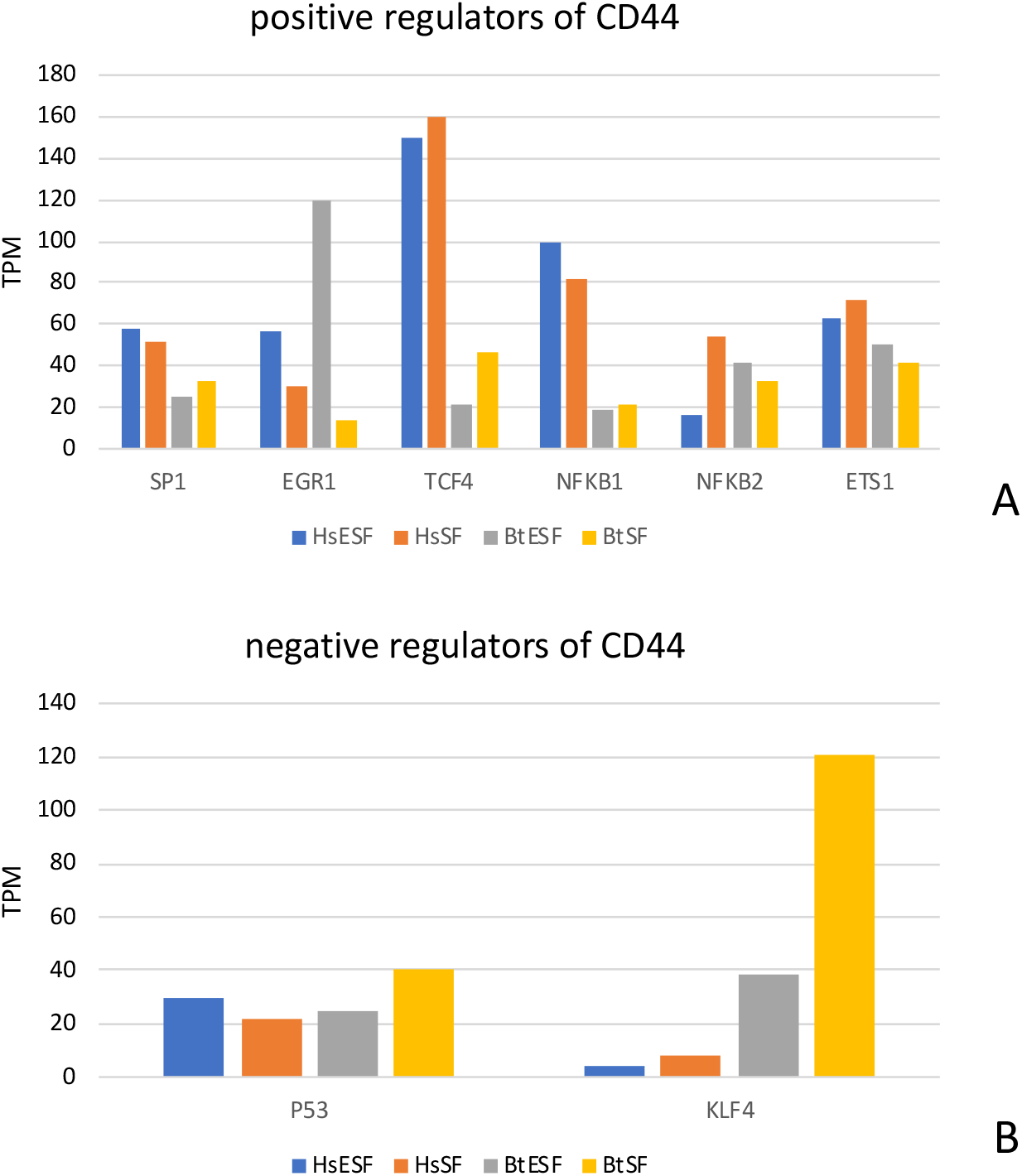
transcription factors known to be regulators of *CD44* in cancer cells. A) RNA expression levels of transcription factors known to be positive regulators of *CD44* in cancer cells. B) RNA expression levels of transcription factors known to be negative regulators of *CD44* in cancer cells.

**Suppl. Figure 7:**
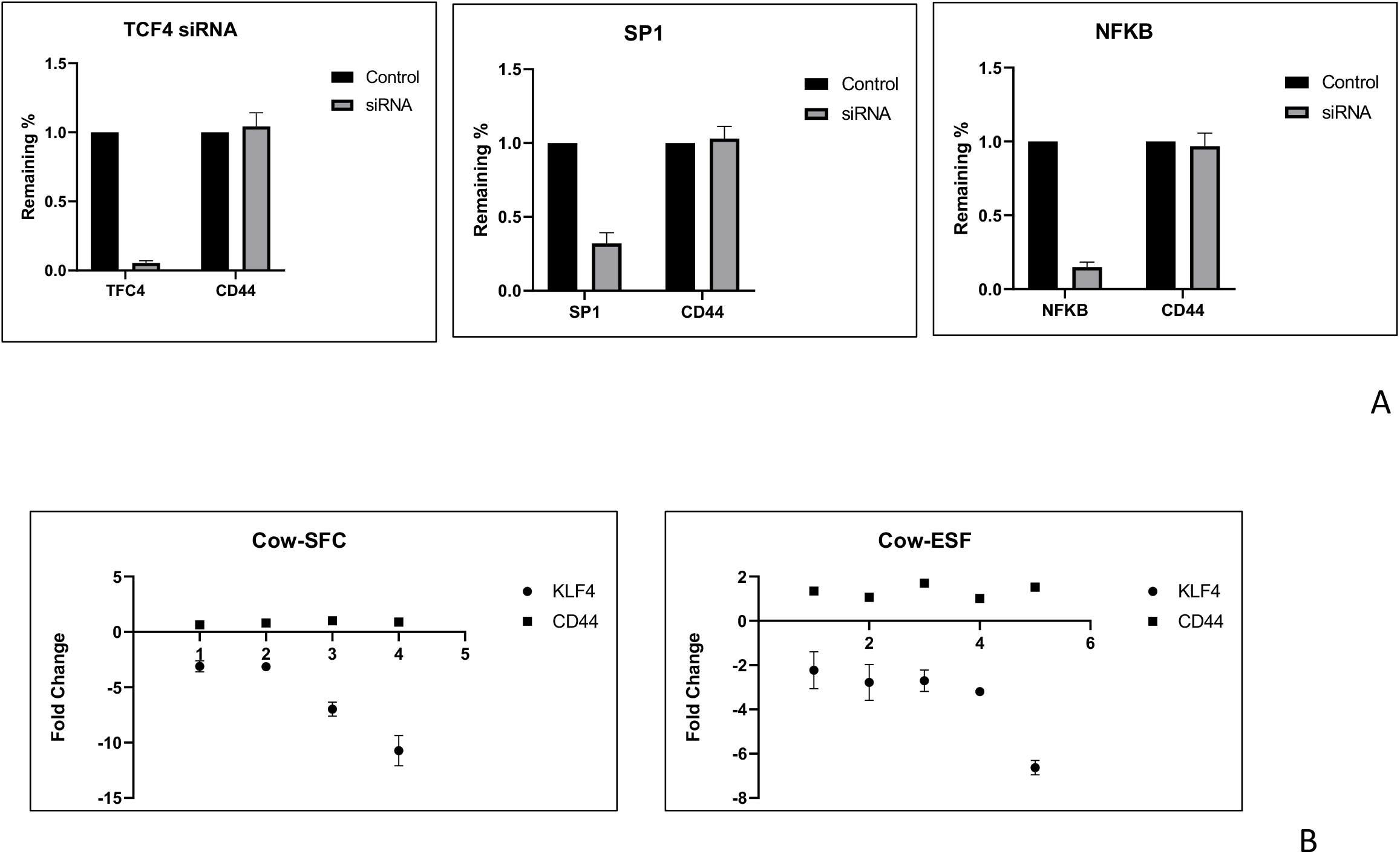
siRNA mediated knockdown of regulators of CD44 which could explain increased expression in humans compared to cow. A) KD results with putative positive regulators of *CD44* in human cells. No effects have been detected. B) KD results with putative negative regulators of *CD44* in cow cells. No effects have been detected.

**Suppl. Figure 8:**
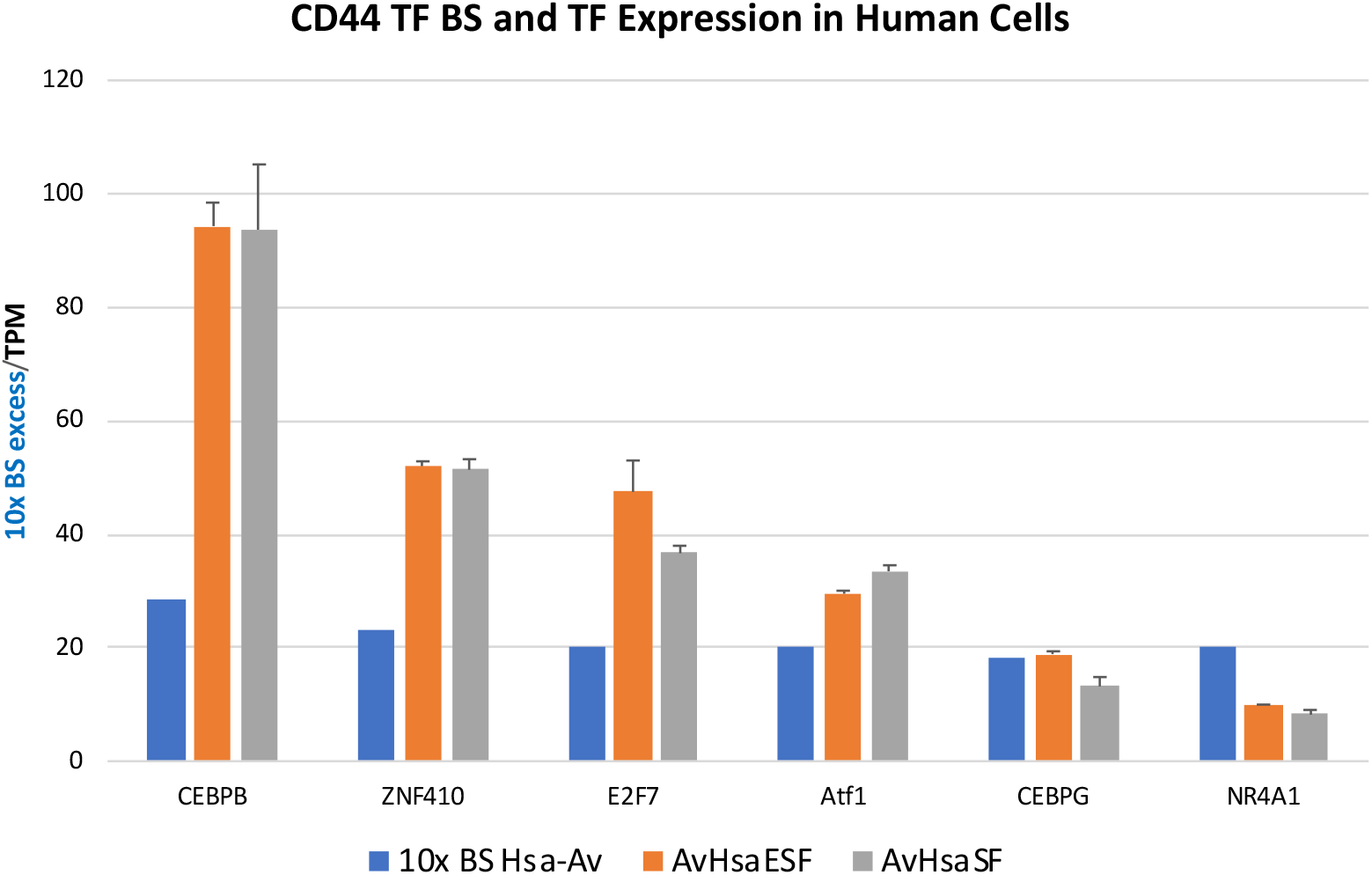
abundance of binding sites and expression levels of cognate transcription factors in human ESF and SF. Blue bars are 10x number of binding sites at the human CD44 promoter, red bar expression level of TF [TPM] in human ESF and gray bar expression level of TF [TPM] in human SF.

**Suppl. Figure 9:**
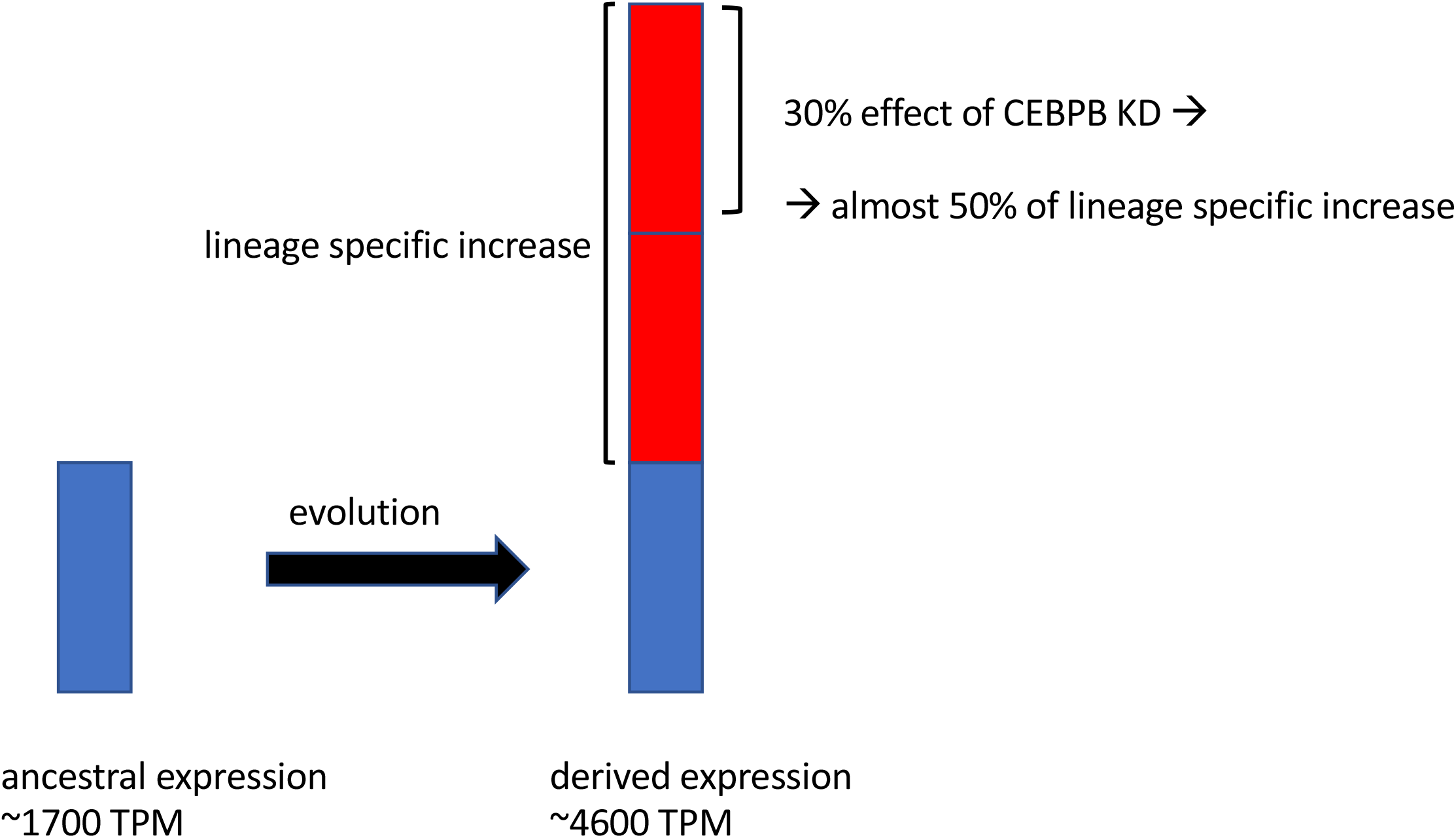
Interpretation of the results in Figure 5A: based on the transcription factor binding site data we think that CEBPB is potentially responsible, in part, for the lineage specific increase of CD44 expression in humans. This increase is about 2.7x, according to the ancestral state reconstruction in Figure 1B. Consequently a 30% reduction of the expression in the derived species (human) accounts for about 50% of the lineage specific increase in CD44 RNA expression.

**Suppl. Figure 10:**
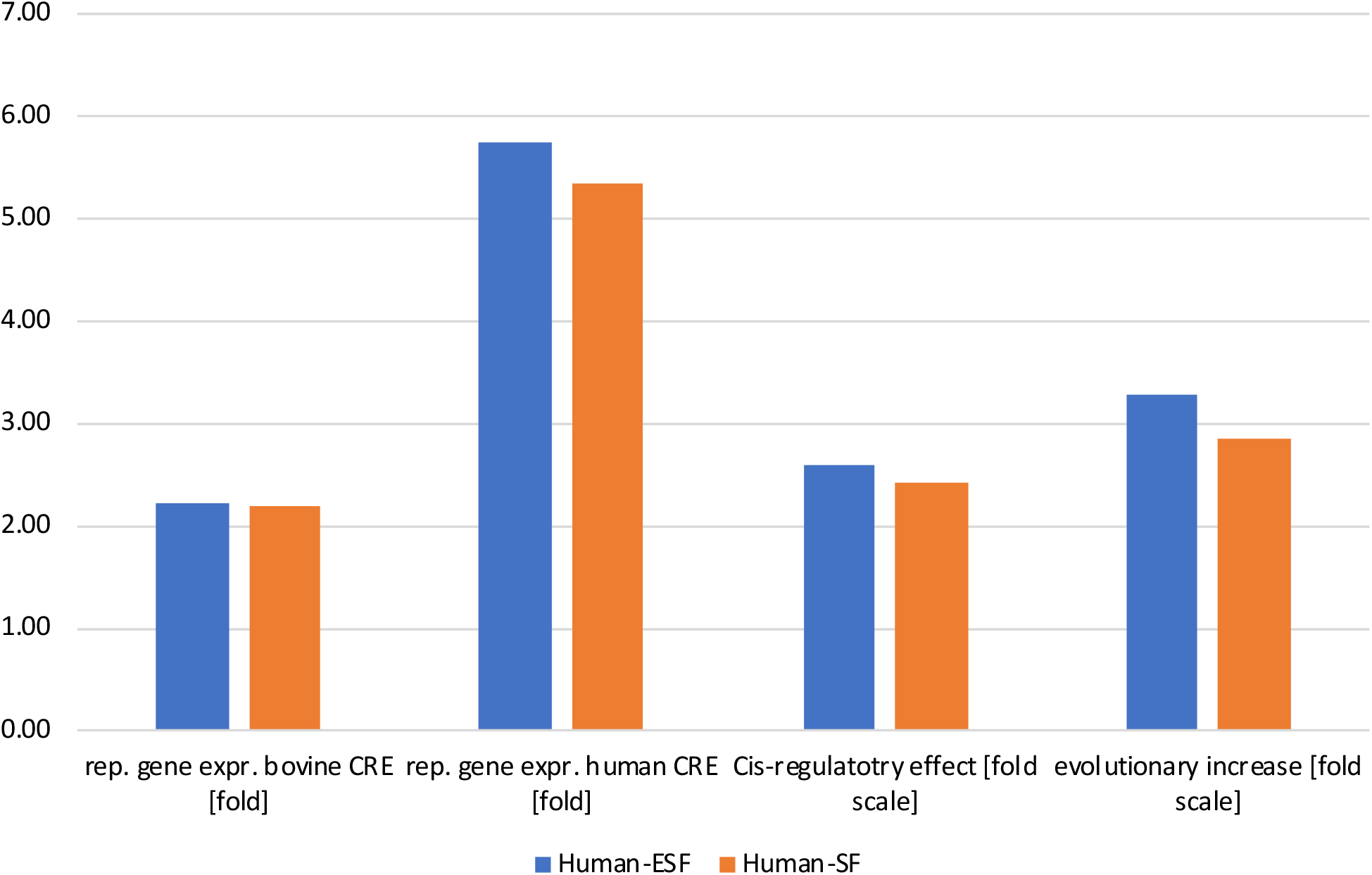
graphical representation of the fold changes in Table 3. Left columns, reporter gene expression driven by the bovine proximal CRE in human cell types, fold difference relative to TBP. Second set of columns, reporter gene expression driven by human CRE in human cell types, fold difference relative to TBP. Third set of columns, cis regulatory effect on the fold scale of replacing the bovine with the human CRE. Right columns, evolutionary change on the fold scale between the inferred ancestral state and the human native CD44 expression.

